# The protooncogene Ski regulates the neuron-glia switch during development of the mammalian cerebral cortex

**DOI:** 10.1101/2022.12.16.520470

**Authors:** Alice Grison, Zahra Karimaddini, Jeremie Breda, Tanzila Mukhtar, Marcelo Boareto, Katja Eschbach, Christian Beisel, Dagmar Iber, Erik van Nimwegen, Verdon Taylor, Suzana Atanasoski

**Affiliations:** Department of Biomedicine, University of Basel, Basel, Switzerland; Department of Biosystems Science and Engineering, ETH Zurich, Basel, Switzerland; Swiss Institute of Bioinformatics (SIB), Basel, Switzerland; Biozentrum, University of Basel, Basel, Switzerland; University Medicine Zurich, University of Zurich, Zurich, Switzerland

## Abstract

The brain is the most complex organ in mammals and understanding the origin of this complexity is a major challenge for developmental biologists. Crucial to the size and morphology of the cortex is the timing and transition of neural stem cell (NSC) fate. An interesting candidate for modulating and fine tuning these processes is the transcriptional regulator Ski, a protooncogene expressed in cortical cells. Ski is involved in diverse cellular processes and epigenetic programs, and mice deficient in Ski exhibit complex central nervous system defects that resemble some of the features observed in patients with 1p36 deletion syndrome and Shprintzen–Goldberg syndrome. Here, we took advantage of *in vivo* transgenic labeling and next-generation sequencing to analyze the gene expression profiles of NSCs, basal progenitor (BP) cells, and newborn neurons (NBNs) from wildtype and Ski-deficient embryos throughout cortical development. We created a unique database that allowed us to identify and compare signaling pathways and transcriptional networks within each progenitor population in the presence and absence of Ski. We find that NSCs are the most affected cell population and uncover that mutant NSCs fail to switch to a gliogenic fate in time. We show that Ski functions in concert with the Bone Morphogenetic Protein (BMP) signaling pathway to alter the cell differentiation fate of NSCs from neurons to glia, which is key to generating adequate numbers of specific cell types during corticogenesis. Thus, by combining genetic tools and bioinformatic analysis, our work not only deepens the knowledge of how Ski functions in the brain, but also provides an immense resource for studying neurodevelopmental disorders.

## INTRODUCTION

In the developing mammalian neocortex, the generation of neurons and glial cells is the result of a balanced proliferative and differentiative division of neural stem and progenitor cells. Regulation of NSC number is associated with rigorous control, and failure of this regulation may lead, for example, to brain malformations, learning and memory disorders, or tumor development. In the initial phase of cortical development, NSCs expand their pool by symmetric proliferative divisions in the ventricular zone (VZ) of the dorsal forebrain. Later, they additionally produce BPs that constitute the adjacent telencephalon-specific subventricular zone (SVZ). The heterogeneous pool of neural stem and progenitor cells sequentially generate molecularly and functionally distinct projection neurons, before perinatally starting to form astrocytes and oligodendrocytes of the neocortex [1–5]. Concomitant cessation of neurogenesis and onset of gliogenesis is achieved by repression of neurogenic and activation of pro-gliogenic genes in the NSC population [6–9]. The use of RNA sequencing technologies, including in our own recent study [10], has enabled detailed analysis of the transcriptomic signature of different cell populations in the developing cerebral cortex [11–16]. These studies now provide a comprehensive basis for exploring the transcriptional landscape of cortical cell populations in neurodevelopmental disorders, as the signaling pathways that are perturbed in central nervous system diseases are the same pathways that play a role in normal development.

The transcriptional regulator Ski interacts with numerous intracellular proteins, thereby modulating gene regulation, signaling pathways, and epigenetic programs [17, 18]. However, hardly any of these functions have been tested under physiological conditions in vivo. In one of the few studies, we have shown that Ski enables the interaction of the chromatin-remodeling and DNA-binding protein Satb2 with the NuRD/HDAC complex in callosal neurons, which in turn initiates crucial processes for their specification during normal cortical development [19]. Ski-deficient mice (Ski*^−/−^*) display complex phenotypes, such as defects in the morphogenesis of craniofacial structures and in the development of the nervous system. Ski*^−/−^* pups die at birth, and their phenotypes resemble some features of individuals diagnosed with 1p36 deletion syndrome. Since the human SKI gene has been mapped to 1p36, loss of Ski activity in these patients may contribute to some of the observed phenotypes [20, 21]. SKI has also been identified as causative gene for Shprintzen–Goldberg syndrome, an autosomal dominant disorder with neurological anomalies [22, 23].

NSCs express Ski at high levels throughout development, while its expression in projection neurons is transient and particularly high in young neurons [49]. We have demonstrated that loss of Ski leads to altered cell cycle properties of neural stem and progenitor cells, and that the timing of neurogenesis is impaired. However, the mechanisms underlying this phenotype remained unexplored [19]. In this work, we combined genetic tools and bioinformatic analysis to directly compare the gene expression profiles of isolated NSCs, BPs, and immature NBNs from wildtype and Ski-deficient embryos, spanning the periods of expansion (embryonic day (E) 11.5), neurogenesis (E12.5-16.5), and gliogenesis (E17.5-E18.5). We find that it is primarily the NSCs population that is affected by the loss of Ski and that the changes in NSC transcriptional profiles are particularly evident at late stages of cortical development. Our data show that mutant NCSs are trapped in a prolonged neurogenic phase and fail to switch to a gliogenic fate in time. We discover that Ski is an important regulator of the transition from the neurogenic to the gliogenic phase and show that it exerts this function in cooperation with the BMP signaling pathway.

## METHODS

### Experimental mouse models

Hes::GFP, Tbr2::GFP, and Ski*^+/−^* transgenic lines were generated as previously described [24–26]. Hes::GFP and Tbr2::GFP mice were mated with Ski*^+/−^* mice to generate Hes::GFP Ski*^+/−^* and Tbr2::GFP Ski*^+/−^*. Subsequently, the two mouse lines were intercrossed to generate homozygous mutant Hes::GFP Ski*^−/−^* and Tbr2::GFP Ski*^−/−^* embryos. Ski inactivation in mice results in perinatal lethality [20], so no postnatal tissue can be examined. Experiments were done using embryos of either sex. Ski conditional knock-out mice (Ski^fl/fl^) were generated by Mouse Clinical Institute (ICS) by insertion of LoxP sites flanking exon 1 of the Ski gene. Mice were maintained on a 12hr day-night cycle with free access to food and water under specific pathogen-free conditions according to the Swiss Federal regulations and ARRIVE guidelines. All experiments were approved by the Veterinary Office of the Canton of Basel-Stadt and the Ethics Committee and were performed according to the 3R guidelines.

### Tissue preparation and Fluorescence Assisted Cell Sorting (FACS)

Dorsal cortices from embryonic day E11.5 to E18.5 were micro-dissected and dissociated into single cell suspensions using Papain and Ovo-mucoid mix (previously described by [27]. Cells were washed with L15 medium, filtered through a 40 μm cell sieve and sorted by forward and side-scatter for live cells (control) and gated for GFP-negative (wildtype levels) or GFP+ populations with a FACSaria III (BD Biosciences). For Hes5::GFP Ski*^+/+^* and Hes5::GFP Ski*^−/−^*, the bright GFP+ cells (apical progenitor cells, APs) were collected. For Tbr2::GFP Ski*^+/+^* and Tbr2::GFP Ski*^−/−^*, both the bright (basal progenitor cells, BPs) and dim (immature NBNs) GFP+ cells were collected. GFP+ cells were used for RNA isolation and gene expression analysis. For each time point (E11.5 – E18.5), 3-4 biological replicates were generated.

### RNA isolation and RNA sequencing

Total RNA was isolated from FACSorted GFP+ cells using the TRIzol method (Life Technology). The integrity of the RNA samples was checked using the Agilent 2100 Bioanalyzer. Their concentration was measured using Quant-IT RiboGreen RNA Assays (Life Technologies). Libraries for Illumina sequencing were prepared with the TruSeq RNA Library Prep Kit v2 (Illumina) and checked for quality using the Fragment Analyzer high-sensitivity NGS kit (AATI). SR50 sequencing was performed on Illumina HiSeq2000 and HiSeq2500 systems with v3 and v4 SBS chemistry, respectively. Reads per sample before mapping ranged from 10 to 70 million reads, and most samples had between 10 and 20 million reads.

### Immunohistochemistry, microscopy, and image analysis

Embryos were fixed with 4% paraformaldehyde (w/v), cryoprotected overnight with 30% sucrose (w/v), and embedded in embedding matrix (Leica). 20 μm coronal sections (E12.5 to E18.5) were processed for immunohistochemistry. Sections from embryos younger than E15.5 were prepared from whole embryo heads. Sections were blocked with 5% normal donkey serum in 0.01% Triton X-100 and 0.1M phosphate buffer for 1h at RT. Sections were incubated overnight at room temperature with the primary antibody diluted in a blocking solution of 1.5% normal donkey serum (Jackson ImmunoResearch) and 0.5% Triton X-100 in phosphate buffered saline (PBS). We used antibodies against goat Sox2 (1:200; Santa Cruz Biotechnology sc-17320); rat BrdU (1:400; AbSerotec OPT0030); goat Dcx (1:200; Santa Cruz Biotechnology sc-271390); mouse Gfap (1:500; Sigma G3893); rabbit pHH3 (1:200; Millipore 06-570); chicken GFP (1:500, Aves labs, GFP-1020); rabbit Sox9 (1:200, gift from M. Wegner); rabbit dsRed (1:500, Clonetech Takara 632496 to detect Tomato protein); mouse βTubulin (1:600, Sigma T8660); mouse γTubulin (1:1000, Sigma T6557); rabbit Olig2 (1:500, Chemicon AB9610); rabbit phospho-Smad1/5/8 (1:100, Cell Signaling #95115); rabbit Id1 (1:500, Biocheck BCH-1/#37-2); rabbit Notch (1:300, own production, unpurified); mouse βCat (1:100, BD TransLab 610153). Sections were washed in PBS and incubated for 2 hours at room temperature with the appropriate secondary antibodies in blocking solution. Secondary antibodies (Jackson ImmunoResearch Labs) used were Cy3-conjugated donkey anti-rabbit (1:200), Cy3-conjugated donkey anti-mouse (1:200), Cy3-conjugated donkey anti-goat (1:200), Alexa488-conjugated donkey anti-rat (1:200), Alexa488-conjugated donkey anti-chicken (1:200), Dy647-conjugated donkey anti-mouse (1:200), Dy647-conjugated donkey anti-rabbit (1:200). All cryosections were counterstained with DAPI (5 µg/ml). Sections were embedded in mounting media containing diazabicyclo-octane (DABCO, Sigma) as an anti-fading agent and visualized with fixed photomultiplier settings on a Zeiss LSM510 confocal, Apotome2 microscope, and Zeiss Image Z1 microscope.

### BrdU birthdating

Timed pregnant females received a single i.p. injection of BrdU (50 mg/kg body weight) at E17.5. Pups were collected at E18.5 and processed for BrdU immunohistochemistry. At least three groups of Ski*^+/+^* and Ski*^−/−^* littermates were examined.

### *In utero* electroporation

In utero electroporation was performed in C57Bl/6 pregnant females 13.5 days after vaginal plug detection as previously described [28] and embryos were collected 48hrs later. The shSki GFP (pSUPER_RETRO Neo/GFP shSki, Oligo engine) and shLuc GFP (pSUPER_RETRO Neo/GFP shLuc, Addgene) plasmids were prepared using endotoxin-free conditions. The shSki sequence cloned into the pSUPER_RETRO Neo/GFP vector was 5’-CAGTGTCTGCGAGTGAGAA-3’ (Oligo engine) [29].

### Neurosphere cultures, viral infection, PCR, RT-qPCR, and sphere forming capacity assay

Primary NSCs were isolated from E13.5 dorsal cortices and cultured in DMEM/F12 + Glutamax medium (containing 2% B27 and 20 ng/μl FGF2) as neurospheres [27]. Ski*^+/+^* and Ski^fl/fl^ neurospheres were infected with adeno-Cre-GFP adenovirus (titer 10^12^ infectious particles/ml [30] to assess recombination. 1x10^8^ infectious particles were used to infect 1x10^6^ cells [31]. 48hrs post infection, genomic DNA was extracted, and flox-recombination events were examined by PCR using 5X FirePol (Solis Biodyne).

Primers for testing excision of the Ski floxed allele were:

A: 5’-ACGAGTCCATCTCTCTGACACTCAGCG -3’;

B: 5’-AGACATCAAACACAGACCTGGGCA -3’

F: 5’-GCCAGCACAAATGGGGACAGTAGTAT -3’

For the RT-qPCR experiment, total RNA was isolated from FACSorted Tomato+ Ski*^+/+^* and GFP+ Ski^fl/fl^ neurospheres 48hrs after infection with adeno-Cre-Tomato using the TRIzol method (Life Technologies) and resuspended in water. RNA was treated with RNase-free DNase I (Roche) to remove genomic DNA contamination. First-strand cDNA was generated using BioScript (Bioline) and random hexamer primers, followed by quantitative RT-PCR (RT-qPCR) using the SensiMix SYBR kit (Bioline). Expression analysis of genes of interest was performed on a Rotor-Gene Q (Qiagen).

Primers for RT-qPCR were:

GAPDH: 5’- GATGCCCCCATGTTTGTG -3’; 5’- GTGGTCATGAGCCCTTCC –3’

mSki: 5’- GAAGGAGAAGTTCGACTATGCC -3’; 5’- TGTGAGGAAGTGTCGTCTGTTT-3’

To test the ability to form spheres, Ski*^+/+^* and Ski^fl/fl^ neurospheres were infected with retro-Cre-Tomato retrovirus. Replication-deficient retroviruses were generated by transfecting Platinum-E cells (Cell Biolabs) with pMI-NLS-Cre-ires-Tomato plasmid using the CalPhos kit (Clontech). Retroviral supernatants were harvested after 48hrs, viruses were purified using the Retro-X Concentrator kit (Clontech) according to manufacturer’s instructions, resuspended in TNE buffer, aliquoted, and stored at -80°C. 10μl of virus was used to infect 1x10^6^ cells. Five days after infection, the spheres where dissociated, and the FACSorted Tomato+ cells were seeded in 24 multi-well plates (500 cells per well). Five days after seeding, a defined number of infected neurospheres were dissociated and the number of recombined secondary spheres formed was quantified. The ratio between the initial spheres and secondary spheres formed provided information about the number of NSCs still present in the infected population. Neurospheres were counted in 24-well plates using an Axio microscope.

### Adherent NSC culture, viral infection, nucleofection and immunostaining

Primary NSCs were isolated from E13.5 dorsal cortices and cultured in DMEM/F12 + Glutamax medium (containing 2% B27 and 20 ng/μl FGF2) as adherent culture [28]. Adherent NSCs were infected with retro-Cre-Tomato retrovirus (5μl of virus was used to infect 3x10^5^ cells) or nucleofected with expression constructs according to instructions of the Amaxa nucleofection kit (Agilent) [28]. Expression constructs used were: pMI-NLS-Cre-ires-Tomato, pMI-GFP (control), pMI-Id1-ires-GFP, and pMI-Id3-ires-GFP. Ski*^+/+^* and Ski^fl/fl^ adherent cells were grown in 12 multi-well plates (8x10^5^ cells per well) and nucleofected with 0.625μg of the respective plasmid. Two days after transfection, differentiation was initiated either by removing FGF2 (neuronal differentiation) or by treating with LIF (100ng/ml) twice every two days (glial differentiation). After 10 days of differentiation, cells were fixed with 4% paraformaldehyde (w/v) for 10 minutes at RT, washed and incubated overnight at 4°C with primary antibodies diluted in blocking solution with 10% normal donkey serum, 1% bovine serum albumin (BSA), and 0.2% Triton X-100 in PBS. After washing, cells were incubated for 1hr at RT with secondary antibodies diluted in blocking solution. Stained coverslips were mounted on glass slides (VWR by Thermo Scientific), embedded in mounting medium (DABCO, Sigma), and visualized using a Zeiss Image Z1 microscope.

### Cloning of Id1 and Id3 into pMIG plasmids

The coding sequences of Id1 and Id3 (468 and 360 nucleotides respectively) were cloned into the pMIG plasmid using the restriction enzymes BglII and XhoI. In both constructs, a Kozak consensus sequence (GCCACC) was added upstream of the ATG. Id1 and Id3 coding sequences were subcloned from pDrive Id1 and pMIG Id3 plasmids, respectively (provided by Christian Schachtrup).

### Quantification and data analysis

Images were optimized for size, color, and contrast using Photoshop CS4 (Adobe). At least three image fields per sample were analyzed and quantified. Sample size is given in the Excel spreadsheets for the quantifications. The number of phospho-histone H3 (pHH3)-positive cells was determined as the number of cells per mm ventricular length. We performed all cell quantifications in the medio-lateral extent of the rostral cortex. We distinguished between proliferating cells in M-phase lining the ventricle (apical) and proliferating cells in M-phase distant from the edge of the ventricle (basal). In the analysis of the IUE experiment, quantifications were performed for GFP+ cells to analyze cell autonomous effects. For cleavage plane orientation analysis, we stained sections of Ski*^+/+^* and Ski*^−/−^* brains with DAPI and γ-tubulin, a centrosome marker, to detect cells in the anaphase/telophase of mitosis at the apical surface of the VZ. We calculated the percentage of NSCs with perpendicular or parallel cleavage planes with respect to the ventricular surface from the total number of apical divisions. Percentages were converted by log10 transformation. Statistical comparisons were performed using the two-tailed unpaired Student’s t-test (if not differentially indicated). Statistical significance was assessed using GraphPad Prism software (GraphPad Software) and set at p < 0.05. All figures were created in Adobe Illustrator CS6. Selected, stained cells were analyzed with fixed photomultiplier settings on a Zeiss LSM510.

### Data and software availability

The RNA sequencing datasets have been deposited in the Gene Expression Omnibus (GEO) under accession number GEO: GSE134688 (Ski*^+/+^*) and GSE134690 (Ski*^−/−^*).

For ISMARA, data can be found here: https://ismara.unibas.ch/NeuroStemX_Ski/.

### Reads mapping and data processing

Reads from cell population mRNA-Seq were mapped to the transcriptome (GENCODE Release M2 GRCm38.p2) with kallisto 0.43.0[*]. The option --pseudobam was used to save the pseudoalignments to transcriptome in BAM file. Reads mapping to multiple transcripts were uniformly distributed over these transcripts. To obtain the expression per transcript, we first divided the number of reads mapping to each transcript by the length of the transcript in nucleotides and then renormalized these length-normalized read counts to transcripts per million (TPM). Gene expression levels were obtained by summing the expression levels of the transcripts corresponding to the gene. Promoter expression levels were obtained by summing for each promoter the length-normalized count of the transcripts associated with the promoter and then re-normalizing to TPM. We added a pseudo-count of 0.5 to the transcript, gene and promoter expression levels in logarithmic space (i.e., log (TPM+0.5)). For the population mRNA-Seq, we also computed replicate averages in log (TPM+0.5). The method used was adapted from Bray et al, 2016 [32, 33].

### Differentially expressed gene analysis

Pairwise comparisons between different time points (conditions) have been applied by using tximport and Deseq2 packages in R on the gene level mRNA expression values (estimated counts from Kallisto). log2 fold change of one and adjusted p-value of 0.01 are considered as cut-off to select differentially expressed genes.

### Dimension reduction and visualization

Samples are illustrated in two dimensions using Principal Component Analysis (PCA). PCA has been employed on centered log transformed TPM gene-level estimates (with pseudo-count of 0.5, log2 (transcripts per million+0.5)) using prcomp function in R. To project Ski*^−/−^* samples on wildtype PCs, first the first two principal components of wild-type samples were calculated. Then, the Ski*^−/−^* expression matrix (containing log2 transformed values) was multiplied by these two vectors.

### Functional annotation analysis

Functional annotation analysis was performed by employing MetaCore for Pathway Maps, Process Networks, Diseases (by Biomarkers), GO Process. For Notch, "Signal transduction_NOTCH signaling", for BMP, "Signal Transduction_BMP and GDF signaling", for Wnt, "Signal transduction_WNT signaling", for Shh, "Development_Hedgehog signaling", for TGF-beta, "Signal Transduction_TGF-beta, GDF and Activin signaling", and for ESR, "Signal transduction_ESR1-membrane pathway" terms were used to export their components for Fig. 4a and Ext.Data Fig. 3.

## ISMARA

We performed ISMARA analysis [34] on the transcriptome-wide gene expression levels of all samples to infer transcription factor motif activities and their target genes of each regulatory motif.

### Principal component analysis on TF activity

We performed principal component analysis on the motif activities inferred with ISMARA separately for the Ski*^+/+^* samples (Ext.Data Fig. 2a) and the Ski*^−/−^* samples (Ext.Data Fig. 2c). To compare the Ski*^+/+^* samples with Ski*^−/−^* samples, the motif activities of each Ski*^−/−^* sample were projected on the first two principal components computed from the Ski*^+/+^* samples (Ext.Data Fig. 2e). In order to visualize the most variable motifs along developmental time we identified the 20 motifs with the highest contribution to the first two principal components, clustered them based on their projection on the first two principal components and visualized them as axis on the first 2 principal components (Ext.Data Fig. 2b, d , f) m, with panels (b), (d) and (f) corresponding to panels (a), (c) and (e) respectively.

### Hierarchical clustering

To identify groups of genes that show similar difference in expression between the time courses of the Ski*^+/+^* and Ski*^−/−^* samples we proceeded as follows. For each gene *g* and time point *t* we calculated a z-score *zgt* corresponding to the difference in log-expression values of gene *g* at time point *t* between the Ski*^+/+^* and Ski*^−/−^* samples, divided by an error-bar for this expression difference. Each error bar was calculated as the square-root of the sum of the log-expression variances across replicates and a term reflecting the Poisson noise of the RNA-seq measurement (which is mainly relevant for low expressed genes). We retained only the genes for which |*zgt*| > 3 for at least one time point and then clustered the z-score profiles of these genes using Ward clustering [35]. Based on the clustering scores, we decided to cut-off the Ward clustering when 6 clusters were remaining. Principal component analysis of the z-scores *zgt* shows that the first 3 PCA components capture 83% of the total variance and that the 6 clusters correspond to major axes along which genes are distributed when projected on these first 3 PCA components, supporting the validity of the clustering results (Ex.Data Fig. 2i). For each cluster we then calculated average normalized expression profiles separately for the Ski*^+/+^* and Ski*^−/−^* samples (Fig. 3). In the same way hierarchical clustering was performed on the z-score profiles of the differences in motif activities across the time course of the Ski*^+/+^* and Ski*^−/−^* samples.

**Figure 1.**
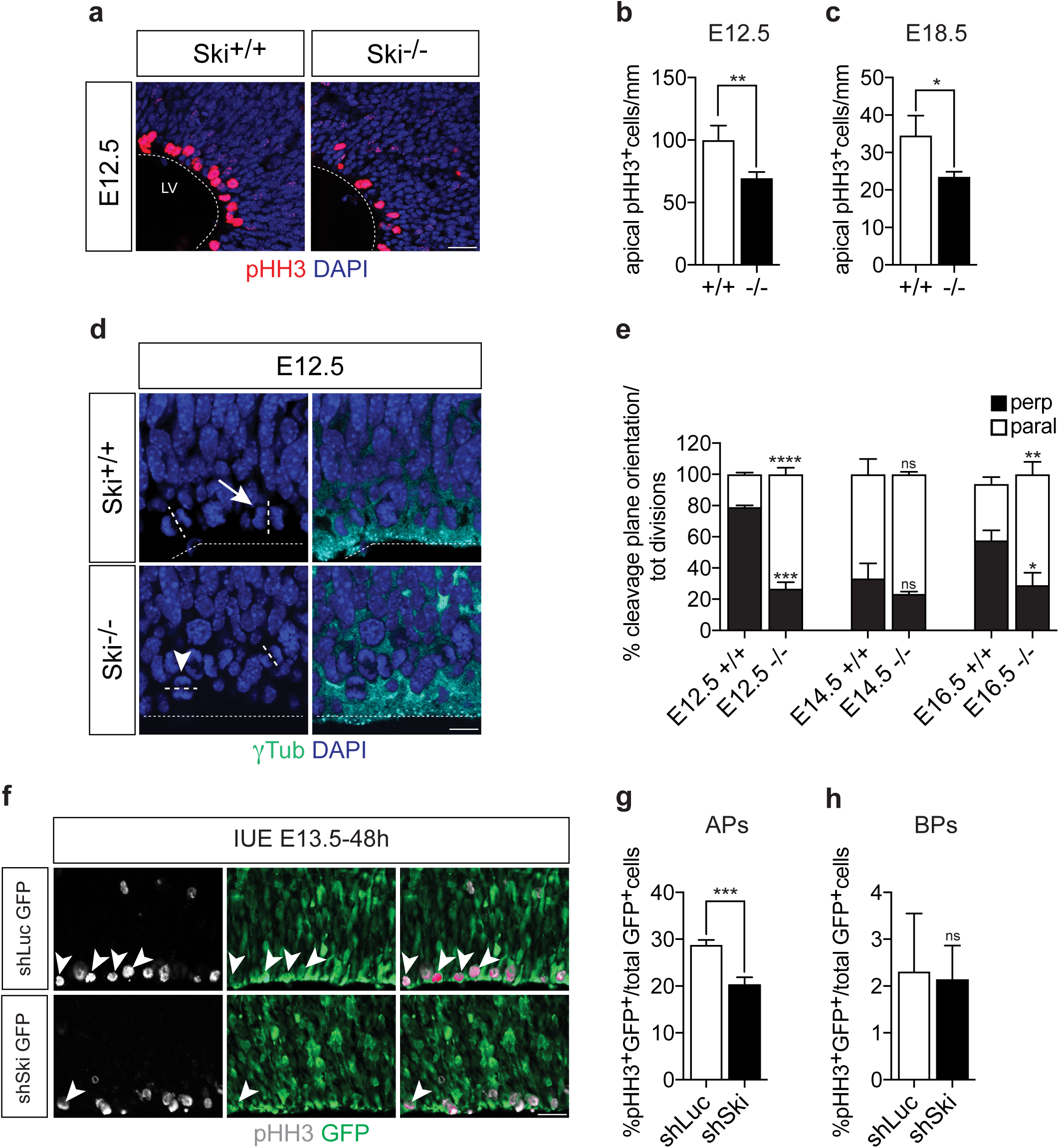
Ski-deficient neural stem cells (NSCs) display proliferation defects throughout development. **(a-c)** Ski*^+/+^* and Ski*^−/−^* NSCs proliferation is analyzed at early (embryonic day E12.5) and late (E18.5) stages of development. **(a)** Immunostainings for phospho-histone H3 (pHH3), a marker for cells in M phase, and nuclear DAPI staining of horizontal E12.5 Ski*^+/+^* and Ski*^−/−^* forebrain sections are shown. **(b, c)** Quantification of the number of pHH3-positive apical NSCs per ventricular length (dashed line in **a**) in Ski*^+/+^* and Ski*^−/−^* sections at E12.5 (**b**) and E18.5 (**c**). Scale bar represents 50µm. LV, lateral ventricle. **(d)** Cleavage plane orientation in Ski*^+/+^* and Ski*^−/−^* apical NSCs is analyzed throughout embryonic development. Immunostaining for γ Tubulin (γTub) and DAPI staining of E12.5 forebrain sections are shown. Dashed lines indicate the cleavage plane orientation, dotted lines mark the apical surface of the lateral ventricle. Scale bar represents 20µm. **(e)** The percentage of perpendicular (perp) (arrow in **d**) or parallel (paral) (arrowhead in **d**) cleavage plane orientation over the total number of mitosis occurring at the apical surface is calculated comparing Ski*^+/+^* and Ski*^−/−^* sections at E12.5, E14.5 and E16.5. In the absence of Ski, an increased proportion of parallel cleavage plane orientated divisions are observed suggesting an increased differentiative division rate at early (E12.5) and late (E16.5) stages. The statistical analysis compares the parallel (white bars) and the perpendicular (black bars) plane orientations between Ski*^+/+^* and Ski*^−/−^* at each time point. (**f-h)** Knock-down of Ski reduces the proliferation of apical progenitor (AP) but not basal progenitor (BP) cells. Proliferation rates are analyzed at E15.5, 48h after *in utero* electroporation (IUE) of shRNA-ires-GFP constructs targeting either Ski (shSki) or Luciferase (shLuc) as control. (**f**) Immunostainings for pHH3 and GFP are shown, arrowheads point to mitotically active GFP-positive cells. Scale bar represents 30µm. (**g**) Quantification of the number of pHH3-GFP double-positive APs (arrowheads in **f**) over the total number of GFP-positive APs in control (shLuc) and shSki-targeted samples. (**h**) Quantification of the number of pHH3-GFP double-positive BPs over the total number of GFP-positive cells per field in control (shLuc) and shSki-targeted samples. At least three experiments were analyzed. *p value= 0.05, **p value=0.01, ***p value=0.001, ns= not significant, t-test was used.

**Figure 2.**
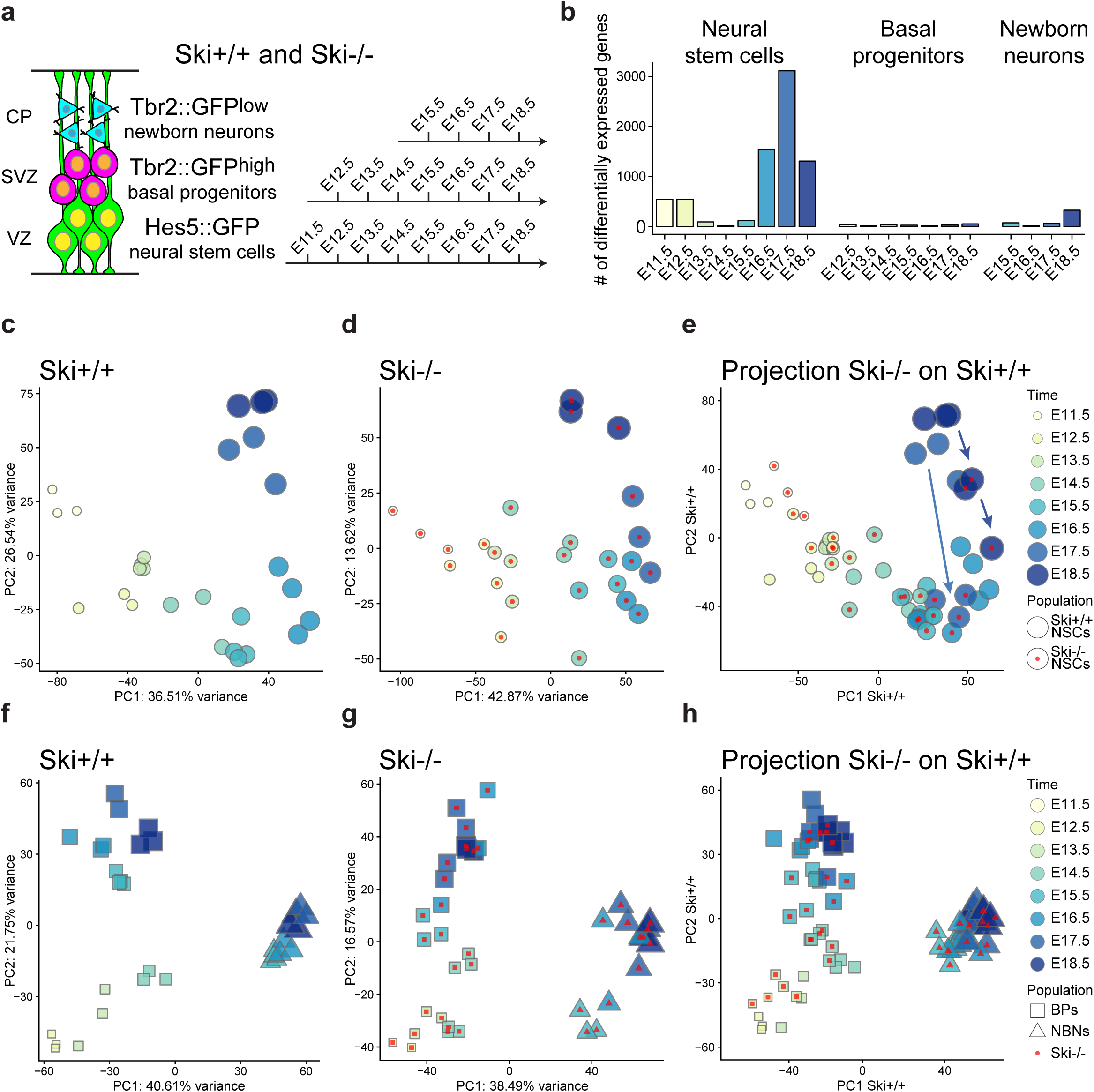
Experimental setup and transcriptional profiles of Ski*^+/+^* and Ski*^−/−^* cortical cell populations over time. **(a)** Schematic representation of the experimental paradigm. Neural stem cells express GFP protein under the *Hes5* promoter (Hes5::GFP), basal progenitors and newborn neurons express GFP protein under the *Eomes* (Tbr2) promoter at high (Tbr2::GFP^high^) and low (Tbr2::GFP^low^) levels, respectively. As indicated, cells of each population were isolated at different days of embryonic (E) development by FACSorting. VZ, ventricular zone; SVZ, subventricular zone; CP, cortical plate. **(b)** The histogram shows the total number of genes differentially expressed when comparing neural stem cells, basal progenitors, and newborn neurons from Ski*^+/+^* and Ski*^−/−^* at each indicated day of cortical development. For the list of genes, see **Table 1. (c-e)** Principal Component Analysis (PCA) of Ski*^+/+^* and Ski*^−/−^* NSCs (c, d). (**e**) Projection of Ski*^−/−^* NSC samples at each time point onto the first two principal components (PC1 and PC2) of Ski*^+/+^* NSCs samples. The arrows point to the affected Ski*^−/−^* NSC samples. (**f-h**) PCA of BPs and newborn neurons (NBNs) using Ski*^+/+^* (f) and Ski*^−/−^* (g) samples. (h) Projection of Ski*^−/−^* BPs and NBNs samples onto the first two PCs of Ski*^+/+^* BPs and NBNs. Shapes without a dot represent Ski*^+/+^* samples and shapes with a red dot represent Ski*^−/−^* samples.

**Figure 3.**
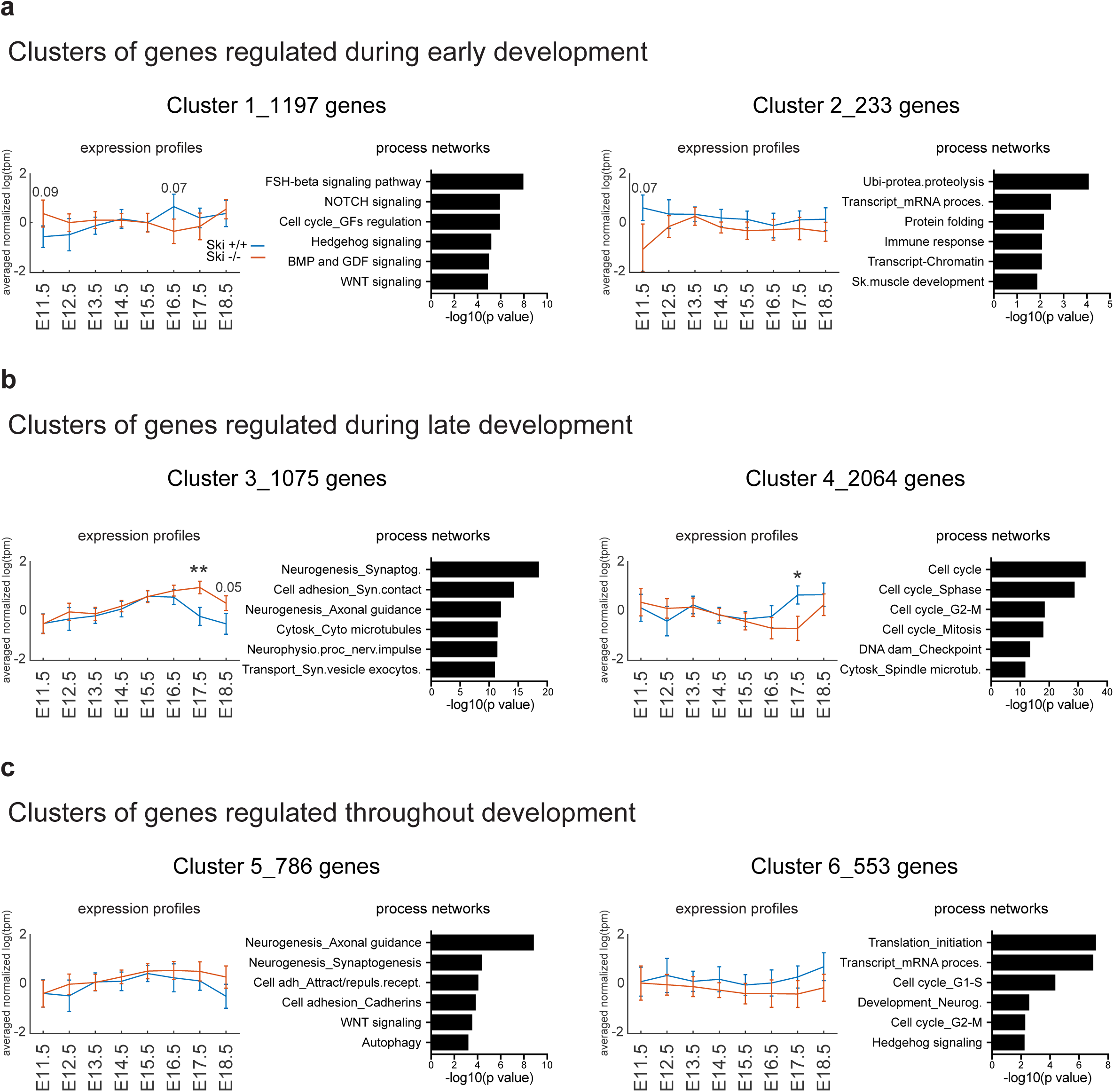
Loss of Ski affects different clusters of genes in NSCs. **(a-c)** Bioinformatic analyses identified 6 clusters of genes differentially expressed comparing Ski*^+/+^* (blue) and Ski*^−/−^* (red) NSCs throughout development. The number of genes in each cluster is indicated. For the list of selected genes see **table2**. Functional annotation analyses show the top 5 process networks enriched in each cluster. **(a)** Clusters 1 and 2 represent genes affected at early stages of corticogenesis (E11.5, E12.5). Genes up-regulated in Ski*^−/−^* NSCs are related to signaling pathways such as Notch1, BMP and WNT (cluster 1). Genes down-regulated in Ski*^−/−^* NSCs are involved in transcriptional and post-transcriptional regulation (cluster 2). **(b)** Clusters 3 and 4 represent genes predominantly affected at late stages (E16.5-E18.5). Genes up-regulated in Ski*^−/−^* NSCs are involved in neurogenesis and neuronal differentiation (cluster 3). Cluster 4 shows genes down-regulated in Ski*^−/−^* NSCs, which are related to the cell cycle machinery. **(c)** Clusters 5 and 6 represent genes differentially regulated throughout development. Cluster 5 is enriched in genes up-regulated in Ski*^−/−^* NSCs at most embryonic time-points. Similar to cluster 3, they are involved in neurogenesis and neuronal differentiation. Cluster 6 shows genes down-regulated in Ski*^−/−^* NSCs, which are related to cell cycle and transcriptional regulation. The expression value is the averaged normalized log tpm of the genes in each cluster. The normalization for each gene is done by subtracting the mean and dividing by the standard deviation. The most significant p values are indicated: *p value= 0.5, **p value=0.01, t-test was used.

**Figure 4.**
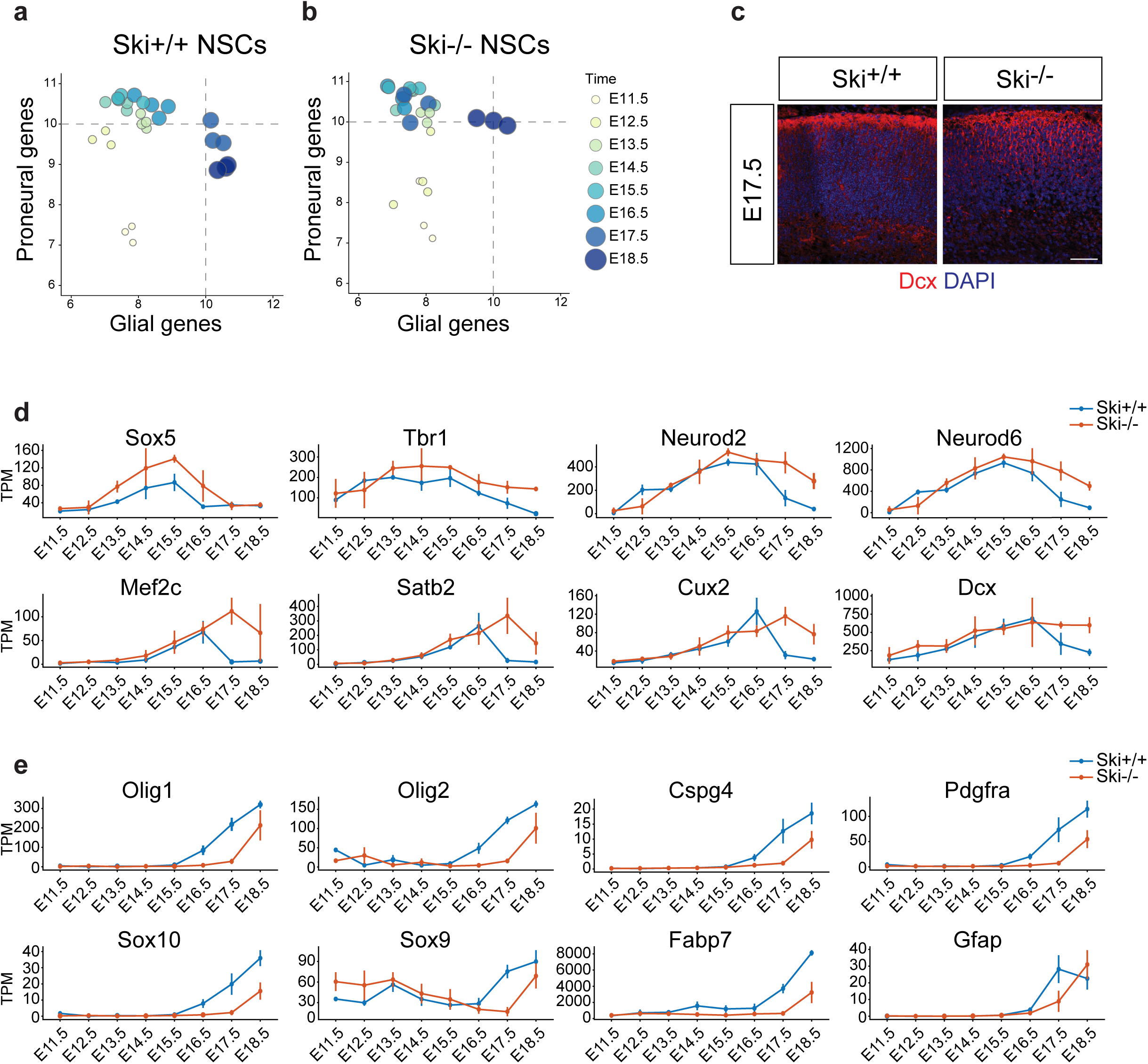
Ski*^−/−^* NSCs persist longer in the neurogenic phase and fail to enter the gliogenic phase in time. **(a, b)** The plots compare total mRNA expression levels in log2 (transcripts per million +0.5) of selected proneural genes (y-axis) and selected glial genes (x-axis) in Ski*^+/+^* NSCs (a) and Ski*^−/−^* NSCs (b). For the list of selected genes see **Table 1**. The intersection of the dashed lines (arbitrary) marks the neuro-glia fate switch in the wildtype. Ski*^−/−^* NSCs still express a high amount of proneural genes at E17.5, while glial gene expression is less efficient than in the wildtype. **(c)** Immunostainings for doublecortin (Dcx), a marker for newborn neurons in the cortical plate (CP), shows a disturbed pattern at E17.5 in the absence of Ski. DAPI, nuclear stain; scale bar represents 50µm. **(d-e)** RNA-seq analysis compares mRNA levels (expressed as TPM, tag per million) of neurogenic (d) and gliogenic (e) markers in Ski*^+/+^* (blue) and in Ski*^−/−^* (red) NSCs. (**d**) mRNA levels of the deep-layer markers Sox5 and Tbr1 and the neurogenic transcription factors Neurod2 and Neurod6 are increased in the absence of Ski. mRNA levels of upper layer markers Mef2c, Satb2, and Cux2 reach the peak of expression in Ski*^+/+^* versus Ski*^−/−^* one day later (at E17.5 instead of E16.5). Moreover, their expression remains at a high level until day E18.5. Dcx mRNA also shows a high expression level at E17.5 and E18.5. (e) mRNA levels of gliogenic markers such as Olig2, Sox9, and Sox10 are downregulated in Ski*^−/−^* NSCs (red) compared to the wildtype (blue) at late stages.

## RESULTS

### Ablation of Ski affects the neural stem cell population in a cell-autonomous manner

Ski protein is expressed throughout the neuroepithelium at E10.5 and is maintained at later stages in the VZ of the dorsal telencephalon, where it colocalizes with NSC markers [19]. Ski*^−/−^* mice show a reduction in radial neuroepithelial thickness and fewer proliferating NSCs at E10.5 and E14.5 [19]. To complement these results, we stained forebrain sections of Ski wildtype (Ski*^+/+^*) and Ski*^−/−^* mice for the mitotic marker phospho-histone H3 (pHH3) and the nuclear stain DAPI at E12.5 (Fig. 1a) and E18.5 and performed mitotic counting. In line with our previous data, we also found a reduced number of proliferating apical NSCs at E12.5 (Fig. 1b) and observed that proliferation defects persisted into late embryonic stages (E18.5) (Fig. 1c).

To extend these results, we next examined and compared the cleavage planes of dividing apical NSCs in Ski*^+/+^* and Ski*^−/−^* cortices (Fig. 1d, e). Apical NSCs undergo predominantly symmetrical proliferative divisions that give rise to two daughter progenitors of the same type as the parent progenitor, and typically require perpendicular (vertical) cleavage plane orientation with respect to the ventricular surface [36]. We stained sections of Ski*^+/+^* and Ski*^−/−^* brains with DAPI and γ-tubulin, a centrosome marker, to detect cells in the anaphase/telophase of mitosis at the apical surface of the VZ (Fig. 1d). Consistent with previous studies, most of the cell divisions that occurred in the wildtype (Ski*^+/+^*) had a perpendicular orientation of the cleavage plane (Fig. 1d, arrow), an effect that was particularly evident in early developmental stages (Fig. 1e, compare Ski*^+/+^* and Ski*^−/−^* black bars at each stage) [37–39]. In contrast, the orientation of the cleavage plane of the dividing apical Ski*^−/−^* NSCs was predominantly oblique or parallel (Fig. 1d, arrowhead) with respect to the ventricular surface throughout development, indicating an increased number of asymmetric cell divisions compared to the Ski*^+/+^* (Fig. 1e, compare Ski*^+/+^* and Ski*^−/−^* white bars at each stage). Asymmetric divisions of NSCs are self-renewing divisions that generate daughter progenitor cells and either other types of progenitor cells, such as BP, or postmitotic neurons [36]. Thus, our results feed into previous observations that the timing of BP production and neurogenesis appears to be disturbed in Ski*^−/−^* mice during cortical development [19].

To differentiate between primary and secondary effects and to investigate whether Ski is directly involved in regulating proliferation of apical NSCs, we introduced an shRNA targeting Ski or a control shRNA (shLuc) together with a GFP vector into E13.5 wildtype brains by *in utero* electroporation and examined the effects on proliferation in apical and basal progenitors 48hrs later (Fig. 1f). We found that Ski inactivation resulted in a significant decrease of pHH3-positive, mitotically active apical NSCs (Fig. 1f, g). Proliferation of BPs was unaffected (Fig. 1h), consistent with the lack of Ski expression in this population [19]. Our results underscore the findings from straight Ski knockout embryos [19, 20] (Fig. 1a-c), and further demonstrate that Ski regulates proliferation of apical NSCs in a cell-autonomous manner. On this basis, we considered the Ski knockout model to be a useful and powerful tool to examine the role of Ski in regulating the dynamics of signaling pathways in NSCs during cortical development and to compare the results with those from other cortical cell populations, such as BPs and NBNs.

### Loss of Ski changes the transcriptional profiles of NSCs particularly at late stages of cortical development

To study and compare changes in gene expression within the different cortical neural lineages of Ski*^+/+^* and Ski*^−/−^* mice, we took advantage of transgenic mice that expressed GFP in a cell-type specific manner (Fig. 2a). Hes5::GFP transgenic mice were previously used for labeling NSCs in the VZ at all developmental stages, while Tbr2::GFP transgenic mice proved suitable for *in vivo* fluorescent labeling of BPs in the SVZ from E12.5, and immature NBNs in the cortical plate from E15.5 on [10]. We show that GFP expression patterns on sections of Hes::GFP Ski*^−/−^* and Tbr2::GFP Ski*^−/−^* embryos were comparable to those of Hes::GFP Ski*^+/+^* and Tbr2::GFP Ski*^+/+^* previously described in detail by Mukhtar et al. (Ex.Data Fig. 1a, b). Cells from individual Hes::GFP Ski*^+/+^*, Tbr2::GFP Ski*^+/+^*, Hes::GFP Ski*^−/−^*, and Tbr2::GFP Ski*^−/−^* embryos were collected and FACSorted at each indicated day of cortical development, as depicted in Fig. 2a. RNA sequencing (RNA-seq) was then performed to gain insight into the transcriptomes of NSCs, BPs, and NBN, and to identify differentially expressed genes in comparison of Ski*^+/+^* and Ski*^−/−^* samples. The results are listed in Table1. Our findings demonstrate that the NSC population is clearly most affected in the absence of Ski, while the BP and NBN population showed little change in the transcriptional profiles between Ski*^+/+^* and Ski*^−/−^* samples (Fig. 2b). This is consistent with the expression of Ski in Sox2 (marker for NSCs) and the absence of Ski expression in Tbr2-positive cells [19]. The largest differences between Ski*^+/+^* and Ski*^−/−^* NSC populations were observed at early (E11.5-E12.5) and late (E16.5-E18.5) time points (Fig. 2b).

To gain greater biological insight into the output of the RNA-seq data, we next performed a functional annotation analysis (Metacore, Pathway Maps) of the genes that were differentially expressed upon deletion of Ski in NSCs (Ext.Data Fig. 1c-g). At early stages (E11.5-E12.5), many gene categories related to signaling pathways, such as Notch, Wnt, Shh, and Tgfβ were found among the first 25 hits (Ex.Data Fig. 1c and Table1), whereas at late stages (E16.5 to E18.5), gene categories related to cell cycle and DNA damage were additionally enriched (Ex.Data Fig. 1d and Table1). We then separately analyzed the genes that were up- or down-regulated in NSCs after deletion of Ski (Ex.Data Fig. 1e and Table1), focusing on the late developmental stages, when NSCs are most affected (Fig. 2b). Among the up-regulated genes, we observed an impressively consistent enrichment of categories related to neurophysiological processes (Ex.Data Fig. 1f and Table1), whereas genes related to the cell cycle machinery, glial differentiation, and various other signaling pathways were down-regulated in the absence of Ski (Ex.Data Fig. 1g and Table1).

We applied Principal Component Analysis (PCA) to discover possible differences in transcriptional dynamics between Ski*^+/+^* and Ski*^−/−^* samples. As recently published, wildtype NSCs display a large variation in gene expression across time on the first two principal components (PCs), and follow a developmental path moving from an expansive to a neurogenic to a gliogenic phase [10] . Interestingly, we uncovered that this transcriptional path was less evident in Ski*^−/−^* NSC samples (Fig. 2d, marked with red dots) compared to Ski*^+/+^* (Fig. 2c). For an even better comparison, we projected the Ski*^−/−^* onto the Ski*^+/+^* PCA (Fig. 2e). Our results showed a clear deviant behavior for the Ski*^−/−^* samples taken at E17.5 and E18.5. They clustered markedly closer to the Ski*^+/+^* samples collected during mid-corticogenesis rather than during late developmental stages (arrows). In contrast, relatively little difference was found between Ski*^+/+^* and Ski*^−/−^* NSCs collected at early or intermediate time points (from E11.5 to E15.5). Thus, PCA illustrates that Ski*^−/−^* NSCs persist in the neurogenic phase and suggest that they do not transition to the gliogenic phase in a timely manner. The analyses of the BP and NBN populations revealed less drastic effects (Fig. 2f-h). The transcriptional dynamics of Ski*^+/+^* (Fig. 2f) and Ski*^−/−^* (Fig. 2g) BPs and immature NBNs were comparable (Fig. 2h), and reflected the RNA-seq data (Fig. 2b).

To characterize the transcriptional state of the Ski*^−/−^* NSCs and map the activities of TFs during cortical development, we used the Integrated System for Motif Activity Response Analysis (ISMARA). ISMARA infers the regulatory states of samples by computationally predicting regulatory sites for TFs throughout the genome. It models the genome-wide expression of each sample in terms of computationally predicted TF binding motifs and their ‘activities’ [34]. PCA of TF binding motifs in Ski*^+/+^* NSC samples showed that, consistent with observations made based on mRNA expression, a specific pool of TFs mediates the developmental path from an expansive to a neurogenic to a gliogenic phase (compare Fig. 2c and Ex.Data Fig. 2a) [10]. We identified the main TF binding motifs in the Ski*^+/+^* and Ski*^−/−^* NSC samples and projected their motif activity vectors onto the same plane to visualize which TFs contribute most to gene expression at each of the developmental stages (Ext.Data Fig. 2b, d, https://ismara.unibas.ch/NeuroStemX_Ski/). To better highlight the differences between the wildtype and mutant NSCs, we projected the Ski*^−/−^* onto the Ski*^+/+^* PCA to jointly analyze the motif activity dynamics (Ext.Data Fig. 2f). This allowed us to define the key TFs and motifs that separate the two genotypes. The motif cluster analysis illustrated very clearly that the major difference in TF activities in Ski*^+/+^* and Ski*^−/−^* NSCs occurred between E17.5 and E18.5 (Ex.Data Fig. 2g). Indeed, some TFs thought to regulate gliogenic transition in Ski*^+/+^*, such as Smad3 and Nfic_Nfib, were not identified in the Ski*^−/−^* NSC samples. In essence, two clusters can be distinguished: the first contains TF motifs that display increased activity in the absence of Ski, such as neuronal repressor Rest [40], the second decreased activity, such as the E2f family involved in cell cycle regulation [41], or Nfia controlling the onset of gliogenesis [42] (Ext.Data Fig. 2h; for the complete list of TF motifs, see Table 2).

### Loss of Ski affects different clusters of genes in NSCs

Unbiased computational analyses revealed extensive transcriptional dynamics between wildtype and mutant NSCs. By hierarchical clustering of gene expression profiles [35], we identified six clusters of differentially expressed genes that follow similar transcriptional trajectories in Ski*^−/−^* relative to Ski*^+/+^* NSCs over time (Ext.Data Fig. 2i and Fig. 3). To determine the six clusters in the hierarchical clustering of the z-score between each Ski*^+/+^* and Ski*^−/−^* sample (see methods), we first performed PCA on the z-score of all genes and all motifs with at least one time point with a z-score of at least |3| (Ext.Data Fig. 2i). We found that 83% of the variance is explained by the first three principal components and that the variance in these three PCs is mainly explained by the z-scores of time points E11.5, E17.5 and E18.5 (see projected axis in Ext.Data Fig. 2i). While the z-score of genes expression separates into six clusters that are positively and negatively aligned with these three axis, the majority of the z-scores of motif activity (84%) are located in only 2 clusters, namely the positive and negative E17.5 clusters (Ext.Data Fig. 2i, red and orange clusters, and Ext.Data Fig. 2g, h). Clusters number 1 and 2 contain genes affected in the early stages of cortical development (Fig. 3a). Even though the RNA levels in the Ski*^+/+^* (blue) did not show significant differences (p value > 0.05) compared to the Ski*^−/−^* samples (red), a tendential difference in the early phases was clearly visible. Cluster 1 is enriched in genes that are upregulated in the absence of Ski at E11.5. Functional annotation analysis (Metacore, process network, see Table 2) revealed that many of them belong to signaling pathways such as Notch, HH, Gdf and BMP (Fig. 3a). Interestingly, these genes are also affected at late time points (E16.5-E17.5), but with an opposite trend in expression. Cluster 2 contains genes that are down-regulated in Ski-deficient NSCs at early developmental stages. However, this cluster is not strongly represented (only 233 transcripts) (Fig. 3a). Cluster 3 and 4 together are the most prominent and significantly regulated clusters with a total of more than 3000 genes up- or downregulated at late stages in Ski*^−/−^* NSCs (Fig. 3b). The upregulated genes belong mainly to categories related to neurogenesis and neurophysiological processes, while the down-regulated are enriched in categories related to cell cycle regulation. The last two clusters 5 and 6 contain genes that are either up- or downregulated throughout development, and similarly to clusters 3 and 4, contain genes related to neurogenesis and cell cycle regulation, respectively (Fig. 3c) (Metacore, process network, see Table 2). Overall, the analysis of the RNA-seq data clearly demonstrates that loss of Ski particularly influences the transcriptional landscape of the NSC population and primarily affects the late stages of cortical development.

### Loss of Ski affects the expression of genes involved in signaling pathways important for cortical development

Functional annotation analysis (Ex.Data 1 c-g) and hierarchical clustering (Fig. 3a) revealed that several genes belonging to particular signaling pathways were affected by the absence of Ski at different time points in NSCs. We therefore analyzed in more detail the transcriptional profiles of modulators and effector targets of some of the key signaling pathways, such as Notch, BMP, TGFβ, estrogen receptor (ESR), and Hedgehog (HH) over time (Ex.Data Fig. 3a). A whole range of differentially expressed genes were found, and remarkably, the early (E11.5, E12.5) and late (E16.5-E18.5) stages were most affected in all selected pathways (Ex.Data Fig. 3a). Staining for Notch1 showed disturbed expression patterns in the absence of Ski at early (Ex.Data Fig. 3b) and late (Ex.Data Fig. 3c) stages. Similarly, the expression of β catenin (βcat), a major component of the WNT signaling pathway, was disturbed at protein level during early (Ex.Data Fig. 3d) and late (Ex.Data Fig. 3e) corticogenesis.

### Ski*^−/−^* NSCs are locked in a prolonged neurogenic phase and fail to enter the gliogenic phase in time

RNA-seq and bioinformatics analyses (Figs. 2, 3) demonstrate that the absence of Ski perturbs the transcriptional landscape of NSCs, especially at late stages of corticogenesis (E16.5-E18.5) when they progress through neurogenesis and start to generate glial cells [1]. We selected a pool of proneural and gliogenic genes to model the behavior of Ski*^+/+^* and Ski*^−/−^* NSCs throughout development (Table 1 and Fig. 4 a, b). We found that Ski*^+/+^* NSCs express many proneural genes (y-axis) during early and mid-corticogenesis (E13.5-E16.5) and then start to express gliogenic genes around E17.5 (x-axis, crossing the vertical dashed line that represents the neuro-glia fate switch) and downregulate proneural genes (y-axis, below the horizontal dashed line) (Fig. 4a). Differently, at E17.5, Ski*^−/−^* NSCs still express high levels of proneural genes, and only at E18.5 do they start to increase gliogenic gene expression to a small extent (x-axis, crossing the vertical dashed line). However, they continue to express proneural genes at high levels (above the horizontal dashed line), even at late stages (E18.5) when repressed in the wildtype (Fig. 4b). Staining for doublecortin (Dcx), a marker for newborn neurons, shows an altered expression pattern when comparing Ski*^+/+^* and Ski*^−/−^* sections at E17.5 (Fig. 4c), providing further evidence that the neurogenic process is compromised.

Comparing the expression profiles of a large number of neuronal markers, we observed that they are consistently upregulated in Ski*^−/−^* NSCs compared to Ski*^+/+^* (Fig. 4d and Ex.Data Fig. 4a). For example, mRNA levels of the deep layer markers Sox5 and Tbr1 and the neurogenic transcription factors Neurod2 and Neurod6 are increased in the absence of Ski. Upper layer markers such as Mef2c, Satb2, and Cux2 reach their peak expression one day later (at E17.5 instead of E16.5) in Ski*^−/−^* compared to Ski*^+/+^*. However, their expression reaches higher levels and remains at high level until day E18.5. A similar profile is seen for Dcx mRNA (Fig. 4d). Conversely, mRNA levels of gliogenic markers such as Olig2, Sox9 and Sox10 etc., are downregulated in late-stage Ski*^−/−^* NSCs (Fig. 4e and Ex.Data Fig. 4b). Taken together, these data show that Ski*^−/−^* NSCs undergo a prolonged neurogenic phase and glial cells do not start to form in time.

### Gliogenesis is impaired in the absence of Ski

To investigate the impaired switch between neurogenic and gliogenic fate in Ski*^−/−^* brains in more detail, we performed a series of experiments to characterize the glial lineage upon loss of Ski. First, we examined gliogenic markers such as Sox9, Olig2, and the glial fibrillary acid protein (Gfap), which are expressed by NSCs late in development, corresponding to the onset of gliogenesis [7]. Immunohistochemical analysis showed that Sox9, an HGM-box transcription factor that is required to specify a glial identity [43, 44], was strongly downregulated in Sox2-positive Ski*^−/−^* compared to Ski*^+/+^* NSCs at E18.5 (Fig. 5a, b). Similar findings were observed for Olig2, a marker for glial progenitor cells (Fig. 5c), and for the astrocytic marker Gfap (Fig. 5d). Next, we assessed the proliferative activity of Ski*^−/−^* gliogenic NSCs. For this, we injected BrdU into pregnant Ski*^+/+^* and Ski*^−/−^* mice at E17.5 and collected embryos 24hrs later. Sections were triple stained for BrdU, Sox2, and Sox9 to identify mitotically active gliogenic NSCs (Ex.Data Fig. 5a). We observed decreased numbers of cells positive for BrdU+Sox2+Sox9+ in Ski*^−/−^* vs. Ski*^+/+^* cortices (Ex. Data Fig. 5b, c), supporting the hypothesis that the gliogenic lineage is affected in the absence of Ski.

**Figure 5.**
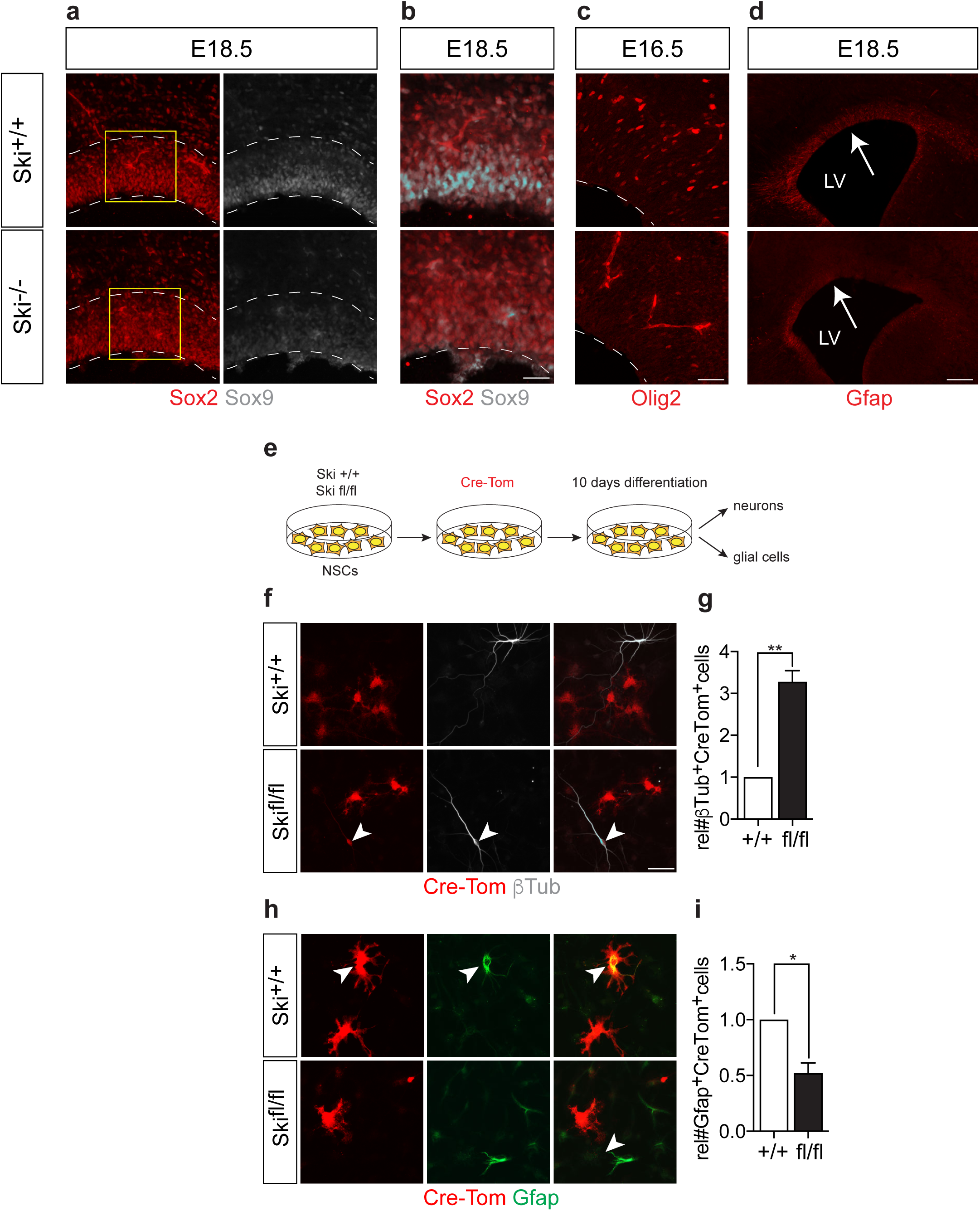
Gliogenesis is impaired in the absence of Ski. **(a, b)** Immunostainings show that the expression of the gliogenic marker Sox9 (white) is strongly downregulated in the ventricular zone (VZ) of Ski*^−/−^* compared to Ski*^+/+^* at E18.5. The dotted lines frame the VZ with the Sox2-positive NSCs (red). Scale bar represents 50 µm. Magnifications of the areas indicated by yellow squares in (a) are shown as merged images of Sox9 and Sox2 double stainings in (b). Scale bar represents 50µm. **(c)** The immunostaining shows fewer cells expressing the gliogenic marker Olig2 in Ski*^−/−^* VZ compared to Ski*^+/+^* at E16.5. The dotted lines indicate the apical surface of the lateral ventricle. Scale bar represents 50µm. **(d)** The immunostaining shows fewer cells expressing the glial marker Gfap in sections of the dorsal cortex of Ski*^−/−^* versus Ski*^+/+^* at E18.5. Scale bar represents 150µm. LV, lateral ventricle. **(e)** Schematic representation of the *in vitro* experiments performed with Ski*^+/+^* and Ski^fl/fl^ NSCs. The cells grown as adhesion cultures were infected with the Cre-Tomato (Cre-Tom) retrovirus. 48h after infection, differentiation was induced by removing growth factors from the media. After 10 days of differentiation, the fate of the infected cells was analyzed. **(f, i)** In the absence of Ski, cultured NCSs differentiate to a greater extent into neurons than into glial cells. **(f)** Immunostainings were performed to detect Cre-Tom-positive cells (red) that were positive for the β Tubulin (βTub) neurogenic marker (white). Arrows point to Cre-infected, βTub-positive cells. Scale bar represents 50µm. **(g)** Quantification of the number of Cre-Tom- and βTub-double positive cells over the total number of infected cells (Cre-Tom+) in Ski*^+/+^* and Ski^fl/fl^ NSC cultures. **(h)** Immunostainings were performed to detect Cre-Tom-positive cells (red) that were positive for the Gfap gliogenic marker (green). Arrows point to Cre-infected, Gfap-positive cells. Scale bar represents 50µm. **(i)** Quantification of the number of Cre-Tom- and Gfap-double positive cells over the total number of infected cells (Cre-Tom+) in Ski*^+/+^* and Ski^fl/fl^ NSC cultures. Data are represented as mean ± SD. At least three experiments were analyzed. *p value= 0.05, **p value=0.01, t-test was used.

To investigate the neurogenic and gliogenic potential of NSCs upon loss of Ski, we took advantage of an inducible Ski knock-out system. For this purpose, conditional Ski floxed (Ski^fl/fl^) mice were generated (see Methods) and neurosphere cultures [45] from E13.5 dorsal cortices were used as starting material, where recombination and excision of a conditional Ski allele was tested (Ex.Data Fig. 6). The Ski wildtype (wt) and Ski floxed (fl) alleles including the selected primers for PCR analysis are depicted in Ex.Data Fig. 6a. Recombination of the conditional Ski allele was induced by infecting neurospheres derived from Ski*^+/+^* and Ski^fl/fl^ mice with a Cre-GFP adenovirus [30]. 48hrs after infection, genomic DNA was isolated and analyzed for deletion at the Ski locus (Ex.Data Fig. 6b). The loxP site-mediated recombination led to a 424 bp product corresponding to the deleted floxed Ski allele that appeared exclusively in the DNA of Cre-GFP/Ski^fl/fl^ (Ex.Data Fig. 6b, lanes 8, 9) but not in the DNA of Cre-GFP/Ski*^+/+^* cells (Ex.Data Fig. 6b, lanes 6, 7). Next, we infected neurospheres derived from Ski*^+/+^* and Ski^fl/fl^ mice with a retro Cre-Tom retrovirus, isolated Tomato+ cells 48hrs later by FACSorting, and performed RT-qPCR experiments. mRNA levels of Ski were reduced by half in Cre-Tom/Ski^fl/fl^ compared to Cre-Tom/Ski*^+/+^* samples (Ex.Data Fig. 6c). To probe *in vitro* cellular mechanisms found in Ski*^−/−^* mice, we carried out sphere forming assays with Cre-Tom/Ski*^+/+^* and Cre-Tom/Ski^fl/fl^ neurospheres (Ex.Data Fig. 6d) and tested neurospheres at passages one, two and three (p1-p3) after retroviral infection. Deletion of Ski in Cre-Tom/Ski^fl/fl^ neurospheres resulted in fewer neurospheres at each passage (Ex.Data Fig. 6e) with smaller diameter (Ex.Data Fig. 6f) compared to the Cre-Tom/Ski*^+/+^* controls. Taken together, our *in vitro* results recapitulate the proliferation defects observed *in vivo* in the Ski knock-out mice [19] (Fig. 1) and strongly support the cell-autonomous function of Ski in NSCs proliferation (Fig. 1).

After validating the inducible Ski knock-out system, we used Ski^fl/fl^ and Ski*^+/+^* adherent NSC cultures (see Methods) to perform differentiation experiments and investigate the neurogenic and gliogenic potential of NSCs upon reducing Ski expression levels. 48hrs after infection with Cre-Tomato retrovirus, differentiation of adherent Cre-Tom/Ski*^+/+^* and Cre-Tom/Ski^fl/fl^ NSCs was induced by removing growth factors from the medium. After 10 days of differentiation, the fate of infected cells was analyzed. The experimental setup is shown in Fig. 5e. Cells were stained for βTubulin (βTub) to detect Tomato-labeled Cre-Tom/ Ski*^+/+^* and Cre-Tom/Ski^fl/fl^ NSCs that differentiated into neurons (Fig. 5f), and for Gfap to label Tomato-positive cells that differentiated into glial cells (Fig. 5h). We observed that a reduction in Ski expression resulted in a significantly higher number of NSCs that were prone to differentiate into neurons (βTub+/Tom+ cells) (Fig. 5g) than into glial cells (Gfap+/Tom+ cells) (Fig. 5i). All these data underline our finding that Ski is an important player in the onset of gliogenesis.

### BMP signaling is affected in Ski*^−/−^* NSCs

In the present study, we have identified a whole series of signaling pathways that are mis-regulated in the absence of Ski (Fig. 3 and Ex.Data Fig. 3). We chose to take a closer look at the BMP signaling pathway (Figs. 6, 7), as it is one of the major determinants involved in the neuronal-glial fate switch at late stages of cortical development [7, 46, 47]. BMPs constitute a family of morphogens that perform a variety of roles at different stages of embryonic development: They inhibit proliferation of NSCs and promote neurogenesis at early stages, and they promote astroglial identity at later stages in development. Specifically, BMP signaling inhibits neuronal lineage commitment and promotes astrocyte differentiation in the late embryonic and postnatal periods. This is reflected at the molecular level, in that activation of BMP signaling leads to increased phosphorylation of Smad proteins (particularly Smad1/5/8) [46, 47] and subsequent transcription of target proteins that inhibit neurogenesis, including Id and Hes factors [47]. Concomitantly, BMPs synergize to promote astrocyte differentiation by activating specific promoters, including the gene encoding Gfap [7, 46, 47]. Hence, multiple cues converge to promote gliogenesis. We stained sections of Ski*^+/+^* and Ski*^−/−^* cortices with antibodies against phospho-Smad1/5/8 (Fig. 6a). Quantifications of apical NSCs revealed a lower number of phospho-Smad1/5/8 positive NSCs in Ski*^−/−^* compared to Ski*^+/+^* samples, suggesting that the BMP signaling pathway is less active in the absence of Ski at E17.5 (Fig. 6b). This is also supported by our sequencing data showing that at the mRNA level, several components of the BMP pathway, such as Id1, Id3, Bmp receptor 1a (Bmpr1a), and Smad9, are downregulated at late developmental stages in cells lacking Ski (Fig. 6c). Additional evidence was provided by immunostaining of sections of Ski*^+/+^* and Ski*^−/−^* cortices for Id1 (Fig. 6d), which revealed a reduction of Id1-positive cells in the VZ as a result of Ski ablation (Fig. 6e). Taken together, these data show that Ski cooperates with the BMP pathway in determining the neuronal-glial fate switch during cortical development.

**Figure 6.**
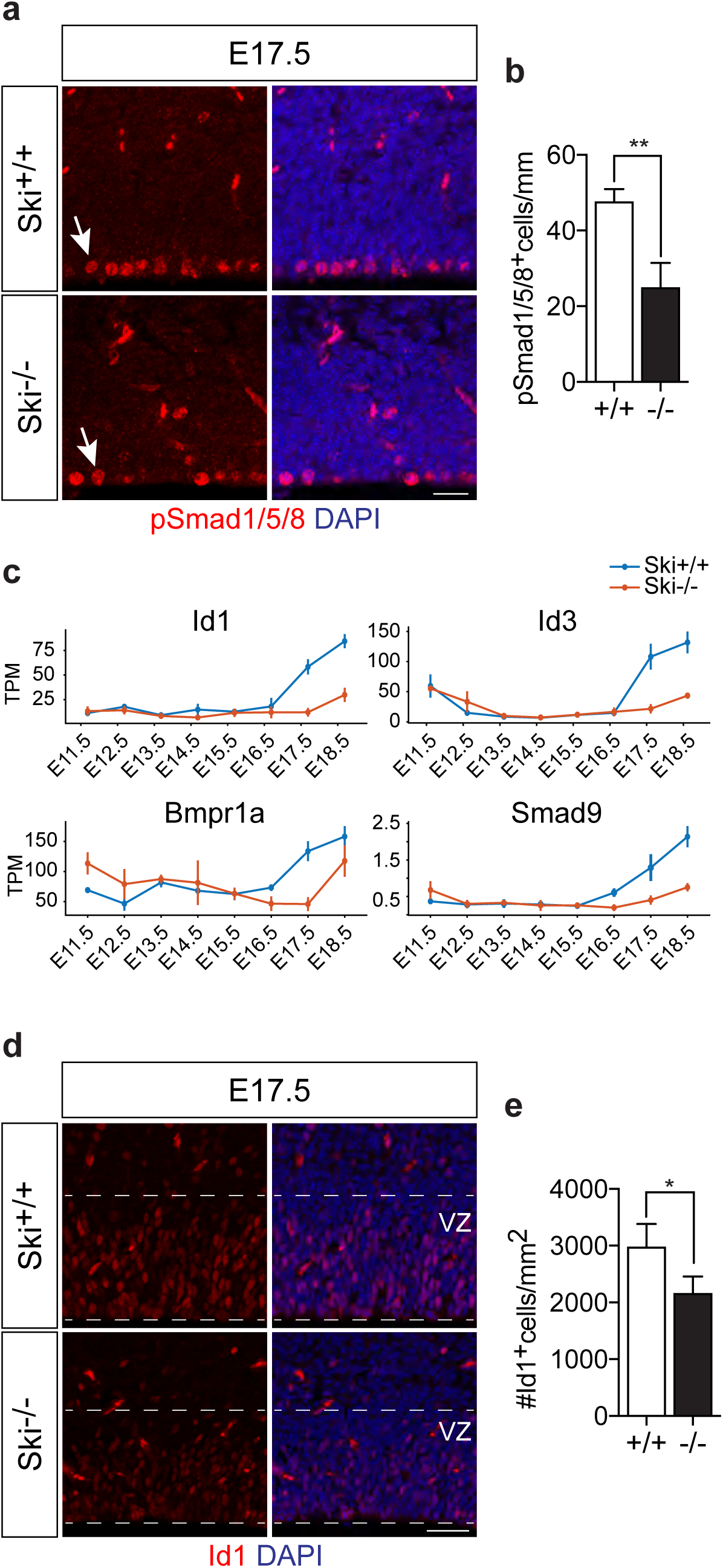
BMP signaling is affected in Ski*^−/−^* NSCs. **(a)** Coronal sections of E17.5 Ski*^+/+^* and Ski*^−/−^* forebrains were immunostained for phospho-Smad1/5/8 (p-Smad1/5/8) and DAPI. Scale bar represents 20µm. **(b)** Quantification shows that ablation of Ski induces a reduction of phospho-Smad1/5/8 positive cells per ventricular length (mm). Only cells lining the ventricle are quantified (indicated by the arrows in (a)). **(c)** RNA-seq analysis shows that components of the BMP pathway, such as Id1, Id3, BMP receptor 1a (Bmpr1a) and Smad9, are down-regulated at the mRNA level (expressed as TPM, tag per million) in Ski*^−/−^* NSCs (red) compared to Ski*^+/+^* (blue) at late stages (E16.5-E18.5). **(d)** Confocal images from Ski*^+/+^* and Ski*^−/−^* coronal brain sections at E17.5 show immunostainings for Id1 in the ventricular zone (VZ). Ski*^−/−^* sections display a reduced number of Id1-positive cells over total DAPI-stained nuclei per field. Scale bar represents 20µm. **(e)** Quantifications show a decreased number of Id1-positive cells per mm^2^ in the VZ in the absence of Ski. Data are represented as mean ± SD. At least three experiments are analyzed. *p value=0.05, t-test was used.

**Figure 7.**
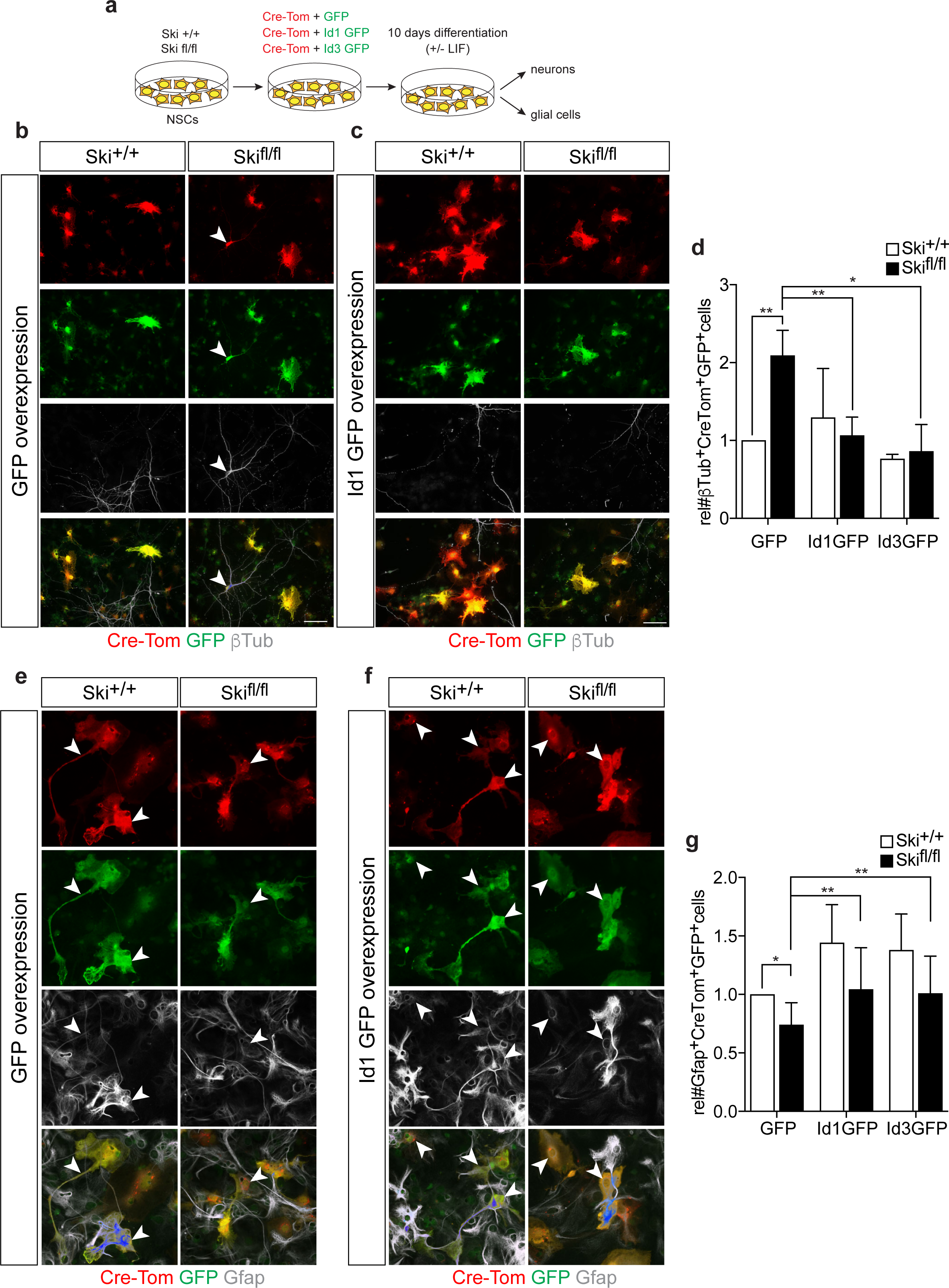
Overexpression of Id1 and Id3 rescues the increase in neuronal differentiation observed upon Ski ablation. **(a)** Schematic representation of the *in vitro* experiments performed with Ski*^+/+^* and Ski^fl/fl^ NSCs. The cells grown as adhesion cultures were transfected with Cre-Tomato (Cre-Tom) plasmid in combination with either Id1-ires-GFP, Id3-ires-GFP or control GFP plasmids. 48hrs after transfection, differentiation was induced by either removing growth factors from the media (b-d) or by removing growth factors and adding LIF to the media (e-g). Following 10 days of differentiation, the neurogenic or gliogenic fate was analyzed by immunostainings. **(b-c)** Immunostainings for Tomato (Cre-Tom, red), GFP (green) and the neuronal marker βTubulin (βTub, white) in Ski*^+/+^* and Ski^fl/fl^ cells upon transfection of Cre-Tom and control (b) or Id1 (c) GFP plasmids. Scale bar represents 50µm. **(d)** Quantifications of the relative number of triple positive cells (indicated by the arrowheads) over the total number of transfected Tom/GFP-double positive cells in cultures derived from Ski*^+/+^* and Ski^fl/fl^ mice. Overexpression of Id1 and Id3 rescues the increase in neuronal differentiation observed upon the reduction of Ski expression levels. **(e-f)** Immunostainings for Tomato (Cre-Tom, red), GFP (green) and the gliogenic marker Gfap (white) in Ski*^+/+^* and Ski^fl/fl^ cells upon transfection of Cre-Tom and control (e) or Id1 (f) GFP plasmids. Scale bar represents 50µm. **(g)** Quantifications of the relative number of triple positive cells (indicated by the arrowheads) over the total number of transfected Tom/GFP-double positive cells in in cultures derived from Ski*^+/+^* and Ski^fl/fl^ mice. Overexpression of Id1 and Id3 partially restores the gliogenic potential impaired in the absence of Ski. Data are represented as mean ± SD. At least three experiments are analyzed. *p value=0.05; **p value=0.01, t-test is used.

### Overexpression of Id1 and Id3 rescues the increase in neurogenesis observed upon Ski deletion

To underpin our results and to further test our hypothesis that Ski functions in cooperation with BMP signaling and Id proteins to regulate the neurogenic-gliogenic fate switch, we performed rescue experiments in conditional Ski-knockout cells. For this, we transfected Ski*^+/+^* and Ski^fl/fl^ adherent NSC cultures with a Cre-ires-Tomato plasmid (Cre-Tom) in combination with either Id1-ires-GFP, Id3-ires-GFP, or GFP (control) plasmid. After transfection, we induced differentiation of the cells for 10 days by removing growth factors from the medium to induce neuronal differentiation, or glial differentiation by also adding LIF (Fig. 7a). Cells were subsequently stained for either the neuronal marker βTubulin (βTub, Fig. 7b, c) or for the glial marker Gfap (Fig. 7e, f) to determine their fate. Quantification of double transfected GFP/Cre-Tom/Ski^fl/fl^ cultures showed a significantly higher number of βTub+ neurons than among double transfected control GFP/Cre-Tom/Ski*^+/+^* cultures (Fig. 7d, GFP, compare white with black bar). These findings were similar but less pronounced than in the experiments using Cre-Tom retroviral infections (Fig. 5g). Overexpression of Id1 did not influence the number of βTub+ neurons in Id1GFP/Cre-Tom/Ski*^+/+^* cultures (Fig. 7d, compare GFP with Id1GFP, white bars). However, it significantly reduced the number of βTub+ neurons in Id1GFP/Cre-Tom/Ski ^fl/fl^ cultures (Fig. 7d, compare GFP with Id1GFP, black bars). Similarly, overexpression of Id3 (Fig. 7d, Id3GFP, black bar) rescued the increased neurogenesis observed upon Ski deletion (Fig. 7d, GFP, black bar), whereas it did not affect the number of βTub+ neurons in control cells (Fig. 7d, compare GFP with Id3GFP, white bars). Regarding Gfap-positive glial cells, quantification of double transfected GFP/Cre-Tom/Ski^fl/fl^ cultures showed a lower number of Gfap-positive glial cells compared to double transfected control GFP/Cre-Tom/Ski*^+/+^* cultures (Fig. 7g, GFP, compare white with black bar). These results, obtained by transfection experiments, were comparable but less pronounced than those obtained by Cre-Tom retroviral infection (Fig. 5i). Overexpression of Id1 or Id3 in Cre-Tom/Ski*^+/+^* cells resulted in enhanced gliogenesis (Fig. 7g, compare GFP with Id1-GFP and Id3-GFP, white bars). Similarly, overexpression of Id1 or Id3 in Cre-Tom/Ski^fl/fl^ cells led to significantly higher numbers of Gfap-positive cells (Fig. 7g, compare GFP with Id1-GFP and Id3-GFP, black bars). However, overexpression of neither Id in Cre-Tom/Ski^fl/fl^ cultures increased the number of Gfap-positive glial cells to the same level as in the control Cre-Tom/Ski*^+/+^* cultures (Fig. 7g, Id1GFP and Id3GFP, compare white with black bars). Hence, overexpression of Id1 and Id3 partially restores the gliogenic potential in Cre-Tom/Ski^fl/fl^ cells, which provides further evidence that Ski is required in the process of gliogenesis. In conclusion, our findings establish Ski as a key modulator of the neurogenic-gliogenic fate switch and show that it exerts this function in cooperation with Id proteins (Fig. 8a).

**Figure 8.**
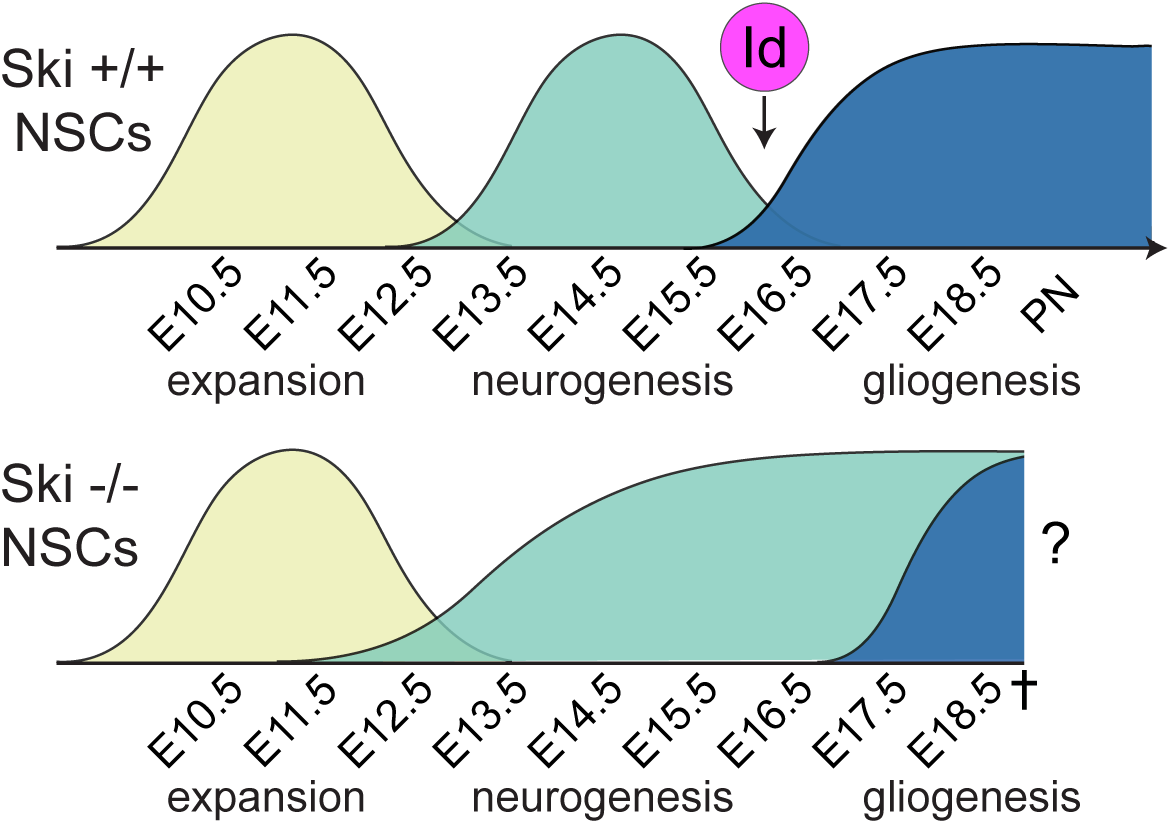
Model for Ski function in NSCs throughout cortical development. In the absence of Ski, the NSCs are trapped in a prolonged neurogenic phase and fail to transition to the gliogenic phase in a timely manner. In the absence of Ski, several signaling pathways are dysregulated, primarily BMP signaling, one of the major determinants of the switch from neuronal to glial phase. We propose that Ski acts as a key modulator of neurogenic-gliogenic fate change and that it exerts this function in cooperation with Id proteins, the main effectors of the BMP pathway.

## DISCUSSION

The protooncogene Ski is linked to various types of cancers as well as to human 1p36 syndrome and Shprintzen–Goldberg syndrome, which are characterized by severe central nervous system defects in addition to other symptoms [18, 20, 22, 23, 26]. Studies performed in normal and cancer cell lines show that Ski is a transcriptional regulator involved in numerous signaling and transcriptional networks, but most of the experiments were performed *in vitro* and using overexpression methods [17, 18]. To date, the knowledge about the physiological function of Ski *in vivo* is limited and discoveries of its molecular modes of action have been made primarily in the nervous system [18, 48, 49]. We have shown that Ski protein has a highly dynamic spatial and temporal expression pattern in the cortex, and its level of expression can strongly influence the molecular machinery of distinct cell types. Ski protein is selectively present in proliferating progenitor cells in the VZ but not in the SVZ, in subtypes of differentiated projection neurons of the cortical plate, but only transiently in newly developing neurons ([19] and Fig. 1). Accordingly, the phenotype of the Ski-deficient cortex is very complex and includes alterations of cell cycle characteristics, premature differentiation of the progenitor pool at early stages of cortical development, and impaired fate specification of callosal projection neurons [19]. In view of the complexity of the system and the highly dynamic nature of Ski expression and regulation, a comprehensive approach such as the one we have taken in this work allows for a rapid overview of cell-type specific and temporal roles of Ski throughout cortical development. The functional diversity of the brain is reflected by its ability to sequentially generate numerous cell types during development. However, it is still unclear how gene expression programs and fate decisions are determined within these different cell populations. In a recent publication, we have presented a unique resource to explore the transcriptional program and the temporal dynamics of gene expression in the developing cortex [10]. This groundwork served as an excellent basis for studying the role of Ski during the expansive, neurogenic, and gliogenic phases of cortical development. Since Ski protein is expressed in Sox2-positive NSCs but not in Tbr2-positive progenitor cells [19], the RNA-seq analyses unsurprisingly revealed perturbations of mRNA expression patterns mainly in NSCs upon loss of Ski. However, the finding that differences were particularly pronounced only at very early and very late embryonic stages, but not at mid-embryonic stages, was not expected. During mid-corticogenesis, NSCs generate BPs from which most cortical neurons will develop. Our previous findings have shown that the BP pool is phenotypically affected by the loss of Ski, as their proliferation rate as well as the number of newly emerging neurons are significantly increased during this time [19]. According to our current findings, the reasons for this tend not to lie in altered mRNA expression patterns of mutant NSCs, BPs, or newborn neurons during the respective time window. Hence, the effects of Ski deletion on the developing cortex appear to involve biological processes that could not be uncovered by our experimental approach but may play out at the protein level. Since Ski protein levels are high in postmitotic neurons [19], transcriptional networks might be impaired in the absence of Ski, leading to perturbed feedback signaling from postmitotic neurons to progenitor cells. This regulatory mechanism, described in previous work, affects the fate specification of progenitor cells (reviewed in [50]) and thus, might also have an impact on their proliferative behavior.

At early developmental stages, the most obvious effect of Ski deficiency is the perturbation of several signaling pathways considered essential during brain development, such as Notch, Hedgehog, and WNT. Our experimental approach also included the analysis of the ligands, receptors, and transcription factors that are part of these signaling pathways. Interestingly, the same signaling pathways are often also affected at late developmental stages, albeit in different ways. That is, a particular gene may show different activity in response to Ski deletion depending on the developmental stage. For example, Notch1 is upregulated at early stages in Ski*^−/−^* compared to wildtype NSCs, but downregulated at late stages, whereas the expression pattern during mid-corticogenesis is unaffected. This result provides insight into a complex scenario in which Ski can activate or repress the transcription of the same gene, possibly depending on whether or not its respective intracellular partners are present. Remarkably, the absence of Ski at early stages virtually invariably leads to upregulation of target genes in all signaling pathways identified, whereas no such unilateral effects are observed at late stages. Ski has historically been considered a transcriptional repressor, but our data imply that Ski may act differently *in vivo* depending on timing and biological context [18, 49]. The stage-dependent differences resulting from Ski deletion are also evident at protein level. In both early and late stages, expression of Notch and ß-catenin in Ski deleted cortices is disturbed, but the pattern differs depending on the day of development.

The strongest effect of Ski deletion in NSCs is seen toward the end of embryonic development when upper layer neurons emerge and the transition from neurogenesis to gliogenesis occurs. Various studies indicate that astrocytes play a critical role in neurodevelopmental diseases such as Rett syndrome and fragile X mental retardation (reviewed in [51]). Hence, perturbation of the timing between neurogenesis and gliogenesis and/or the number of different cells that are generated during embryonic neurogenesis might have a profound impact on the formation and the function of neuronal circuits and result in neurodevelopmental disorders. Although numerous studies suggested that both intrinsic and extrinsic signals interact in setting this complex schedule, the molecular mechanisms are not yet fully understood [7]. In Ski-deficient NSCs, a large group of neurogenic genes show differences in the dynamics of their expression. In support of this *in vivo* data, Ski deletion also causes NSCs to preferentially differentiate towards neuronal lineages *in vitro,* at the expense of glial cell populations. Epigenetic changes in NSCs are thought to be important mechanisms for glial fate switching [52]. Ski could well be an interesting candidate for epigenetic modulation, since it interacts with chromatin-remodeling proteins [18] [49].

BMPs are the main determinants of neuronal-glial fate switch in NSCs and exert a dual function: primarily terminating neurogenesis through the action of the Id (Inhibitor of DNA Binding) proteins and then initiating gliogenesis through the induction of the Gfap gene [7, 46, 47]. The ability of Ski to interact with the BMP pathway has been previously described in Xenopus and in mammalian cells *in vitro* [53], but has not been linked to cortical development. Our experiments provide insight and explanation for some of the features observed in the developing mutant cortex *in vivo* and expand our knowledge of the mechanism of BMP signaling in late cortical development. The next step will be to further decipher the nature of the interaction between Ski and the BMP signaling pathway. This is also of interest in light of the controversy surrounding the interaction of Ski with SMAD proteins. While many reports claim that Ski exclusively associates with TGFβ-specific regulatory Smad2 and Smad3, other groups demonstrate that Ski also interacts with the BMP-Smads and represses BMP signaling [53–55]. However, most of the data showing the specific interplay between Ski and the TGFβ pathway have been obtained in the context of cancer and using *in vitro* overexpression methods. Our new data uncover that the functions of Ski in the physiological context are very broad and its role in BMP signaling is highly relevant. In summary, there is growing evidence that Ski plays an important role in glial biology. Our previous work has demonstrated that Ski is required for myelination in the peripheral nervous system [48], and more recently different studies have established a role for Ski in astrocyte proliferation and migration [56, 57]. In view of this, it will be exciting to further explore the function of Ski in glial biology, during development, in the adult, and under physiological and pathological conditions.

## Supporting information

ExData_Fig_1

ExData_Fig_2

ExData_Fig_3

ExData_Fig_4

ExData_Fig_5

ExData_Fig_6

## Acknowledgements

We thank members of the VT laboratory for helpful discussions, Christian Schachtrup for the generous gift of Id1 and Id3 constructs, and Mikhail Pachkov for designing the website for ISMARA. AG was supported by the University of Basel Research Fund for excellent early career researchers. The senior authors of the NeuroStemX consortium (CB, DI, EvN, VT, and SA) were supported by the Swiss Federal Government (SystemsX.ch –The Swiss Initiative in Systems Biology).

## EXTENDED DATA_FIGURE LEGENDS

**Extended Data Figure 1 Transcriptional profiles of Ski*^+/+^* and Ski*^−/−^* NSCs over time (a, b)** Immunostainings for GFP on cortical sections of Hes5::GFP and Tbr2::GFP Ski*^+/+^* (a) and Ski*^−/−^* (b) embryos at E16.5. Hes5::GFP staining is visible specifically in the VZ, whereas Tbr2::GFP is specifically detectable in the SVZ/IZ. The dotted line delineates the ventricular zone. NSCs, neural stem cells; BPs, basal progenitors; NBNs, newborn neurons; VZ, ventricular zone; SVZ, subventricular zone, IZ, intermediate zone. **(c, d)** Functional annotation analysis (Metacore software, Pathways maps) of genes differentially expressed in Ski*^+/+^* and Ski*^−/−^* NSCs at early (E11.5-E12.5) (c) and late stages (E16.5-E17.5-E18.5) (d). The first 20 enriched categories are shown. Categories of genes related to signaling pathways (early stages) and related to cell cycle regulation (late stages) are particularly detected. **(e)** Histograms show the total number of genes upregulated (grey) or downregulated (green) in Ski*^+/+^* and Ski*^−/−^* NSCs. **(f, g)** Functional annotation analysis (Metacore software, Pathways maps) of genes up-regulated (f) or down-regulated (g) in Ski*^+/+^* and Ski*^−/−^* NSCs at late stages (E16.5-E17.5-E18.5). Among the upregulated genes in Ski*^−/−^* samples, categories related to neurophysiological processes (highlighted in orange) are particularly enriched (f). Among the downregulated genes in Ski*^−/−^* samples, categories related to the cell cycle machinery (highlighted in yellow), glial differentiation (labeled in blue), and signaling pathways (Notch pathway, highlighted in grey) are particularly enriched (g).

**Extended Data Figure 2 Bioinformatic analysis of the transcription factor motif activity in Ski*^+/+^* and Ski*^−/−^* NSCs over time (a-f)** Principal component analysis (PCA) performed on the motif activity of Ski*^+/+^* (a) and Ski*^−/−^* (c) samples. (e) Projection of the Ski*^−/−^* samples on the first 2 components of the Ski*^+/+^* samples, capturing about 80% of the total variance. The color code indicates the sampling time. Right panels (b, d, f): Similar to left panel (a, c, e) with additional projection of the 20 motifs with highest contribution to the first 2 components, grouped in 6 clusters. On (e) and (f), the principal components are computed on the Ski*^+/+^* samples only and the Ski*^−/−^* are projected on those 2 components. On (e) and (f) dots indicate Ski*^+/+^* samples and stars correspond to Ski*^−/−^* samples. **(g)** Difference in motif activity between Ski*^+/+^* and Ski*^−/−^* samples at each time point, relative to the summed standard deviation (Z-score) averaged on the 2 clusters. **(h)** Normalized averaged motif activity in the 2 clusters as a function of the sampling time points for the Ski*^+/+^* sample (blue) and the Ski*^−/−^* samples. The clusters are based on TF activity between Ski*^+/+^* and Ski*^−/−^*. The error bars show the standard deviation. **(i)** Principal component analysis of the z-scores *zgt.* For each gene *g* and time point *t* a z-score *zgt* was calculated corresponding to the difference in log-expression values of gene *g* at time point *t* between the Ski*^+/+^* and Ski*^−/−^* samples, divided by the square root of the sum of the squared error-bar of the the Ski*^+/+^* and Ski*^−/−^* log-expression values. The first three PCA components capture 83% of the total variance and the six clusters (Fig. 3) correspond to major axes along which genes are distributed when projected on these first three PCA components, supporting the validity of the clustering results.

**Extended Data Figure 3 Loss of Ski affects the expression of genes involved in signaling pathways important for cortical development** (**a**) Heatmaps illustrating the expression profile of each gene of selected signaling pathways based on the z-scored log2 (TPM). (**b-e**) Notch1 and β catenin display disturbed patterns of protein expression at early and late stages in the absence of Ski. Immunostainings for Notch1 (b, c) and β catenin (βcat) (d, e) combined with nuclear DAPI staining of horizontal Ski*^+/+^* and Ski*^−/−^* forebrain sections are shown at E12.5 (b, d) and E17.5 (c, e). Dotted lines indicate the apical surface of the ventricular zone. Scale bars represent 20µm.

**Extended Data Figure 4 RNA-seq analysis of Ski*^+/+^* and Ski*^−/−^* NSCs during cortical development (a, b)** RNA-seq analysis comparing mRNA levels (expressed as TPM, tag per million) of a panel of neuronal (a) and glial (b)-related genes in Ski*^+/+^* (dark red) and Ski*^−/−^* (orange) NSCs. The genes analyzed were selected based on published data [58]. At late stages, expression levels of neuronal markers are higher (a), whereas expression levels of glial markers are lower (b) in Ski*^−/−^* compared to Ski*^+/+^*.

**Extended Data Figure 5 Proliferation analysis of Ski*^−/−^* NSCs (a)** Immunostaining for BrdU, Sox2, and Sox9 of E18.5 Ski*^+/+^* and Ski*^−/−^* forebrain sections of embryos injected with BrdU at E17.5. Arrowheads point to triple positive cells. Scale bar represents 50µm. **(b)** The percentage of BrdU, Sox2, and Sox9-triple positive cells (proliferating gliogenic progenitors) over the total number of BrdU and Sox2-double positive cells (proliferating progenitors) in the ventricular zone (VZ) is determined in Ski*^+/+^* and Ski*^−/−^* brain sections. **(c)** The relative number of proliferating gliogenic progenitors (BrdU+Sox2+Sox9+) over the proliferating neurogenic progenitors (BrdU+Sox2+Sox9-) (glia vs neuro index) is reduced in Ski*^−/−^* NSCs. Data are represented as mean ± SD. Three experiments were analyzed. *p value=0.05, **p value=0.01, t-test was used.

**Extended Data Figure 6 Validation of the Ski flox/flox system and proliferation analysis of Ski floxed cells (a)** PCR strategy to test the deletion of Ski in cells derived from Cre-Tom/Ski^fl/fl^ neurospheres with selected primers. A-B product contains the genomic region in which the 5’ loxP site is inserted. This product is 411bp long in the floxed Ski samples. The A-F PCR product is visible only if the floxed exon is excised as a 424 bp band. In the wildtype allele, it would be 12626 bp long. **(b)** Ski*^+/+^* and Ski^fl/fl^ neurospheres were infected with adeno Cre-GFP adenovirus and analyzed after 48hrs. PCR analysis on genomic DNA was performed with selected primers (as indicated in a) to test the deletion of the Ski gene in Ski^fl/fl^ samples upon Cre recombination. Two Ski*^+/+^* samples (lanes 1, 2, 6, 7) and two Ski^fl/fl^ samples (lanes 3, 4, 8, 9) were tested; c (lanes 5, 10) are PCR negative control (H20). **(c)** Relative expression of Ski mRNA level in FACSorted Cre-Tom/ Ski*^+/+^* and Cre-Tom/Ski^fl/fl^ neurospheres. RT-qPCR experiments show that Ski mRNA is reduced upon Cre-recombination in Cre-Tom/Ski^fl/fl^ samples. Data are represented as mean ± SD. Two samples per genotype were analyzed. **(d)** Cre-Tom/Ski^fl/fl^ NSCs display a reduce sphere-forming capacity *in vitro*. Ski^fl/fl^ and Ski*^+/+^* neurospheres were infected with retro Cre-Tom retrovirus, FACSorted after 48 hours and the sphere-forming capacity assays was performed for three weeks. Immunostainings for Tomato (Cre-Tom) and brightfield images are show. Scale bar represents 50µm. **(e)** Quantification of the relative number of spheres formed at each passage (p1, p2, p3) comparing Cre-Tom/Ski*^+/+^* and Cre-Tom/Ski^fl/fl^ neurospheres. **(f)** Quantification of the relative size of Cre-Tom/Ski*^+/+^* and Cre-Tom/Ski^fl/fl^ neurospheres at p1 shows a reduced size of Cre-Tom/Ski^fl/fl^ neurospheres compared to Cre-Tom/Ski*^+/+^* controls. Data are represented as mean ± SD. Three experiments were analyzed. *p value=0.05, **p value=0.01, ***p value=0.001, t-test was used.

## DATA INFORMATION

**Table S1 related to Figure 1.**
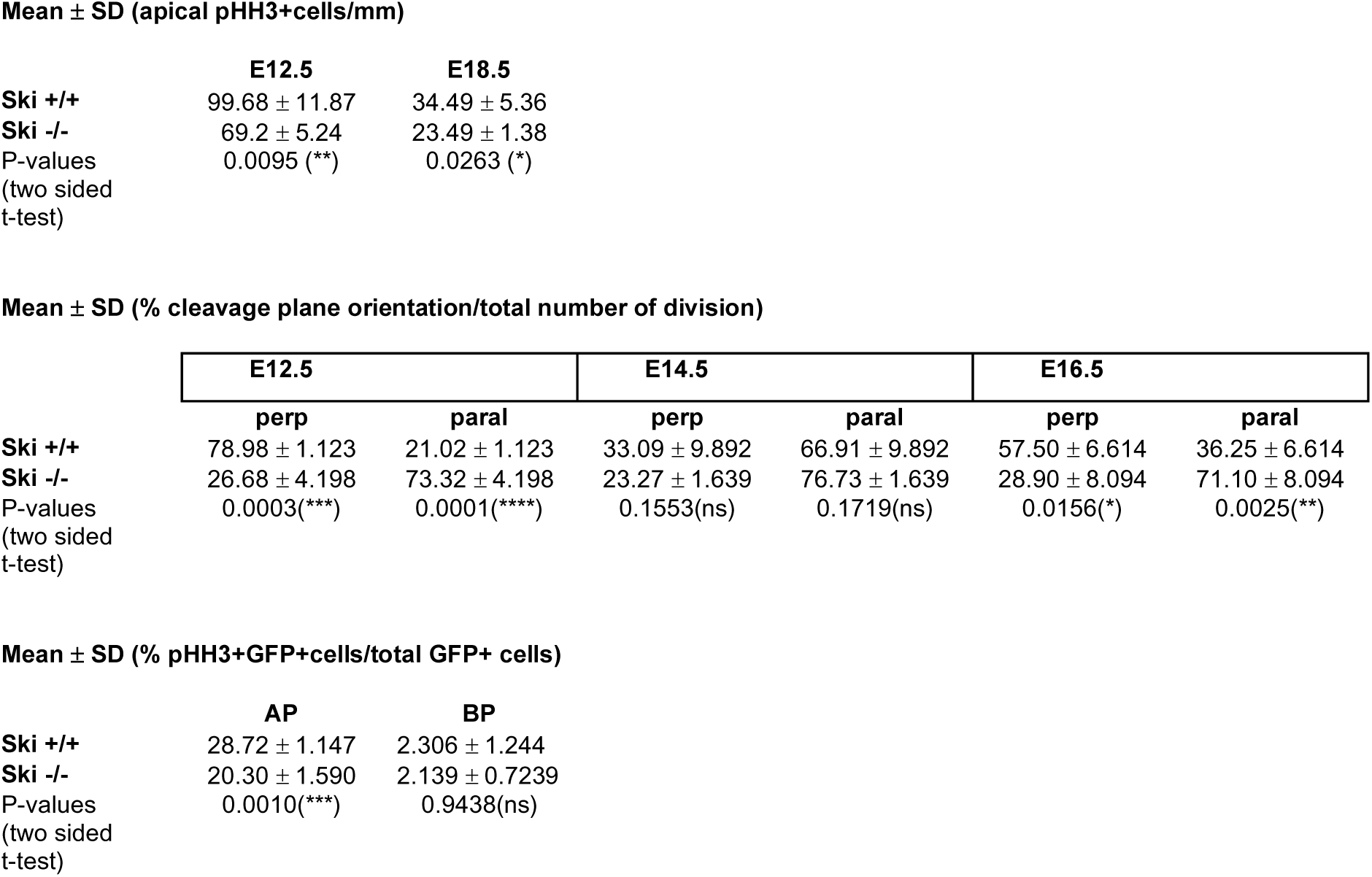

Mean ± SD (apical pHH3+cells/mm)
|  | E12.5 | E18.5 |
| --- | --- | --- |
| **Ski +/+** | 99.68 ± 11.87 | 34.49 ± 5.36 |
| **Ski -/-** | 69.2 ± 5.24 | 23.49 ± 1.38 |
| P-values<br>(two sided<br>t-test) | 0.0095 (**) | 0.0263 (*) |

Mean ± SD (% cleavage plane orientation/total number of division)
|  | E12.5 |  | E14.5 |  | E16.5 |  |
| --- | --- | --- | --- | --- | --- | --- |
|  | perp | paral | perp | paral | perp | paral |
| **Ski +/+** | 78.98 ± 1.123 | 21.02 ± 1.123 | 33.09 ± 9.892 | 66.91 ± 9.892 | 57.50 ± 6.614 | 36.25 ± 6.614 |
| **Ski -/-** | 26.68 ± 4.198 | 73.32 ± 4.198 | 23.27 ± 1.639 | 76.73 ± 1.639 | 28.90 ± 8.094 | 71.10 ± 8.094 |
| P-values<br>(two sided<br>t-test) | 0.0003(***) | 0.0001(****) | 0.1553(ns) | 0.1719(ns) | 0.0156(*) | 0.0025(**) |

**Table S1 related to Figure 1**
|  | AP | BP |
| --- | --- | --- |
| **Ski +/+** | 28.72 ± 1.147 | 2.306 ± 1.244 |
| **Ski -/-** | 20.30 ± 1.590 | 2.139 ± 0.7239 |
| P-values<br>(two sided<br>t-test) | 0.0010(***) | 0.9438(ns) |

**Table S2 related to Figure 5.**

| | βTub | Gfap |
| --- | --- | --- |
| **Ski +/+** | 1 | 1 |
| **Ski fl/fl** | $3.275 \pm 0.27$ | $0.5187 \pm 0.09$ |
| P-values<br>(two sided t-test) | 0.0017(**) | 0.0226(*) |

**Table S3 related to Figure 6.**
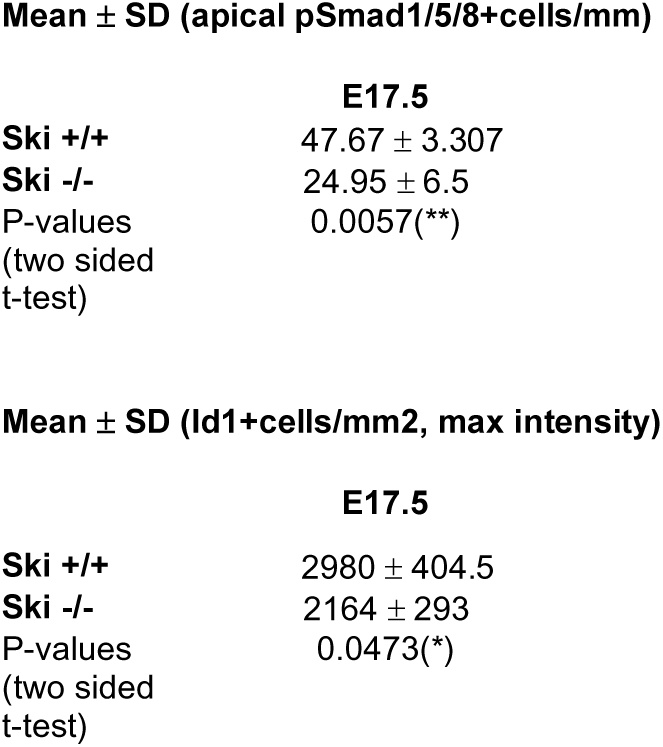

**Mean ± SD (apical pSmad1/5/8+cells/mm)**
|  | **E17.5** |
| --- | --- |
| **Ski +/+** | 47.67 ± 3.307 |
| **Ski -/-** | 24.95 ± 6.5 |
| P-values<br>(two sided<br>t-test) | 0.0057(**) |

**Table S3 related to Figure 6**
|  | **E17.5** |
| --- | --- |
| **Ski +/+** | 2980 ± 404.5 |
| **Ski -/-** | 2164 ± 293 |
| P-values<br>(two sided<br>t-test) | 0.0473(*) |

**Table S4 related to Figure 7.**
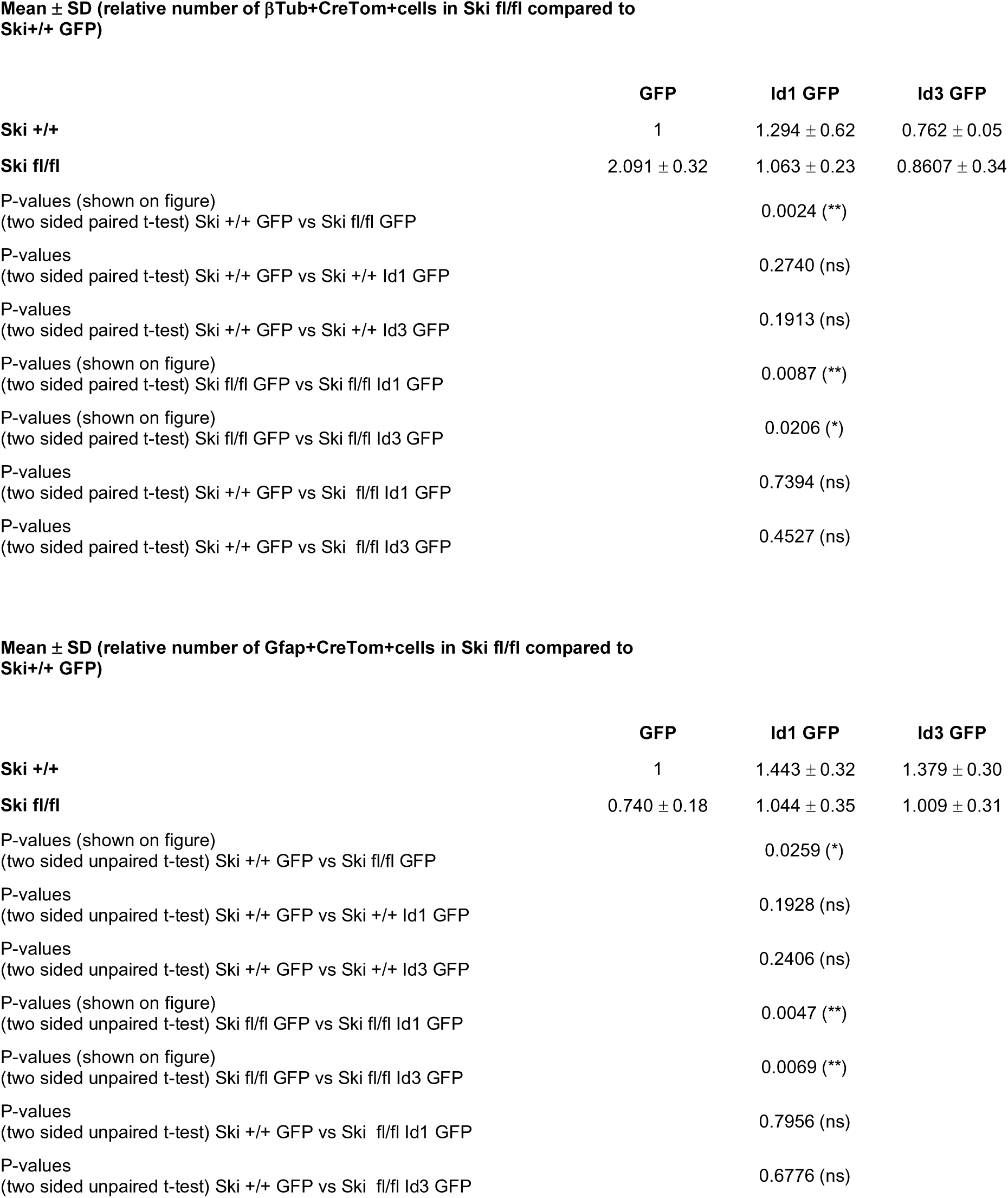

**Mean ± SD (relative number of βTub+CreTom+cells in Ski fl/fl compared to Ski+/+ GFP)**
|  | **GFP** | **Id1 GFP** | **Id3 GFP** |
| --- | --- | --- | --- |
| **Ski +/+** | 1 | 1.294 ± 0.62 | 0.762 ± 0.05 |
| **Ski fl/fl** | 2.091 ± 0.32 | 1.063 ± 0.23 | 0.8607 ± 0.34 |
| P-values (shown on figure)<br>(two sided paired t-test) Ski +/+ GFP vs Ski fl/fl GFP |  | 0.0024 (**) |  |
| P-values<br>(two sided paired t-test) Ski +/+ GFP vs Ski +/+ Id1 GFP |  | 0.2740 (ns) |  |
| P-values<br>(two sided paired t-test) Ski +/+ GFP vs Ski +/+ Id3 GFP |  | 0.1913 (ns) |  |
| P-values (shown on figure)<br>(two sided paired t-test) Ski fl/fl GFP vs Ski fl/fl Id1 GFP |  | 0.0087 (**) |  |
| P-values (shown on figure)<br>(two sided paired t-test) Ski fl/fl GFP vs Ski fl/fl Id3 GFP |  | 0.0206 (*) |  |
| P-values<br>(two sided paired t-test) Ski +/+ GFP vs Ski fl/fl Id1 GFP |  | 0.7394 (ns) |  |
| P-values<br>(two sided paired t-test) Ski +/+ GFP vs Ski fl/fl Id3 GFP |  | 0.4527 (ns) |  |

**Table S4 related to Figure 7**
|  | **GFP** | **Id1 GFP** | **Id3 GFP** |
| --- | --- | --- | --- |
| **Ski +/+** | 1 | 1.443 ± 0.32 | 1.379 ± 0.30 |
| **Ski fl/fl** | 0.740 ± 0.18 | 1.044 ± 0.35 | 1.009 ± 0.31 |
| P-values (shown on figure)<br>(two sided unpaired t-test) Ski +/+ GFP vs Ski fl/fl GFP |  | 0.0259 (*) |  |
| P-values<br>(two sided unpaired t-test) Ski +/+ GFP vs Ski +/+ Id1 GFP |  | 0.1928 (ns) |  |
| P-values<br>(two sided unpaired t-test) Ski +/+ GFP vs Ski +/+ Id3 GFP |  | 0.2406 (ns) |  |
| P-values (shown on figure)<br>(two sided unpaired t-test) Ski fl/fl GFP vs Ski fl/fl Id1 GFP |  | 0.0047 (**) |  |
| P-values (shown on figure)<br>(two sided unpaired t-test) Ski fl/fl GFP vs Ski fl/fl Id3 GFP |  | 0.0069 (**) |  |
| P-values<br>(two sided unpaired t-test) Ski +/+ GFP vs Ski fl/fl Id1 GFP |  | 0.7956 (ns) |  |
| P-values<br>(two sided unpaired t-test) Ski +/+ GFP vs Ski fl/fl Id3 GFP |  | 0.6776 (ns) |  |

**Table S5 related to Extended Data Figure 5.**
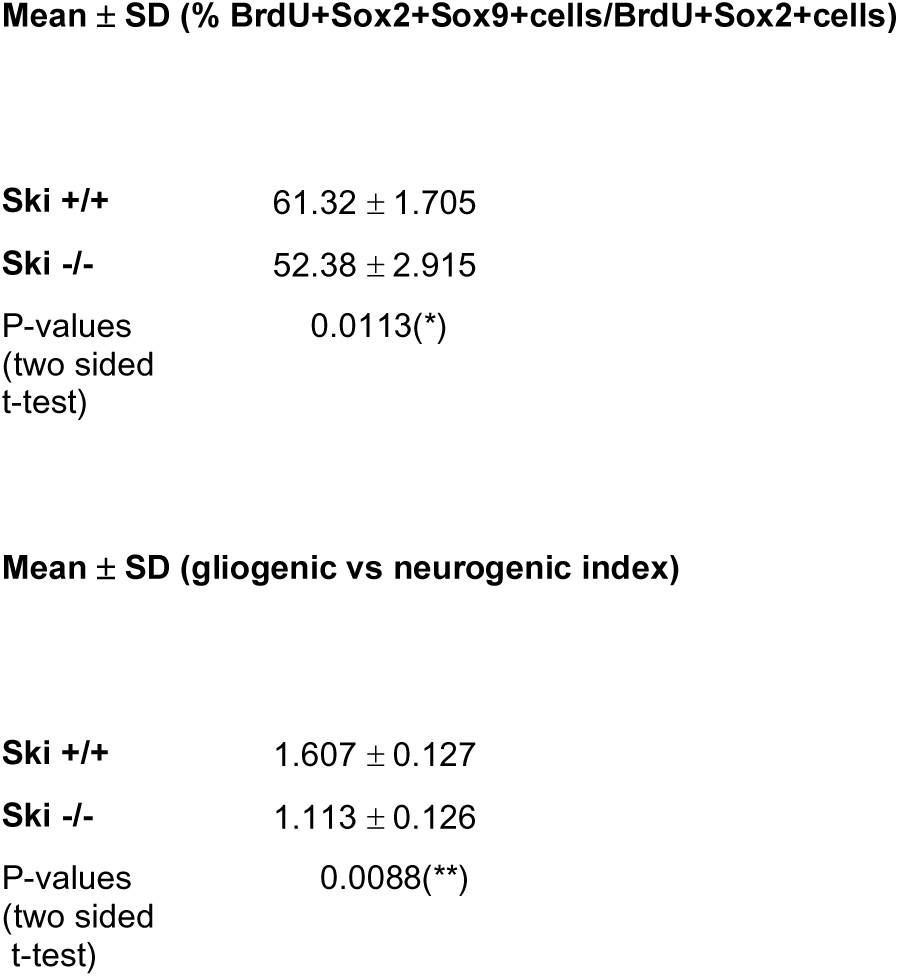

**Table S6 related to Extended Data Figure 6.**
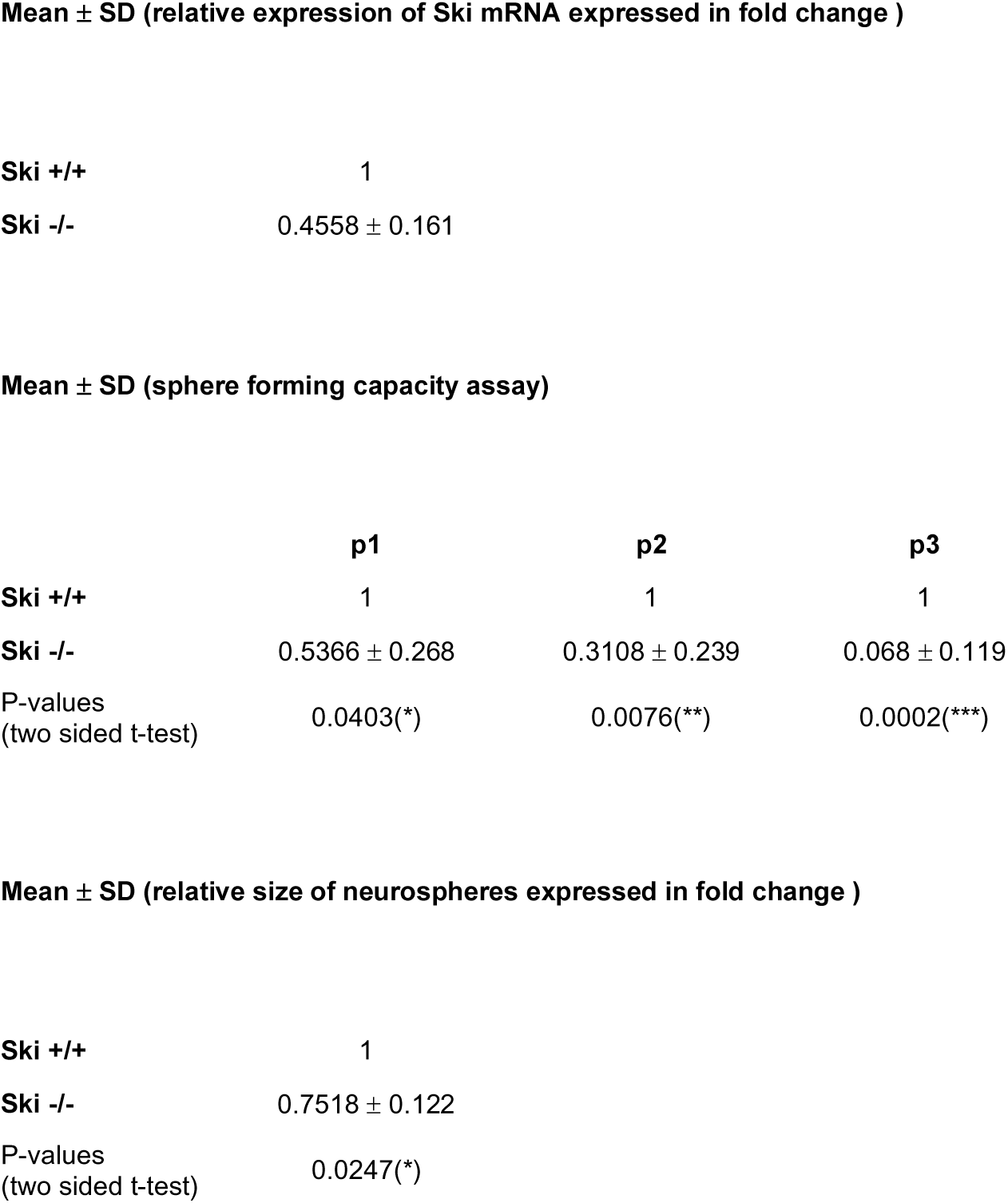

