## Supplementary figures and images for "The protooncogene Ski regulates the neuron-glia switch during development of the mammalian cerebral cortex"

### ExData_Fig_1

Extended Data_Figure1

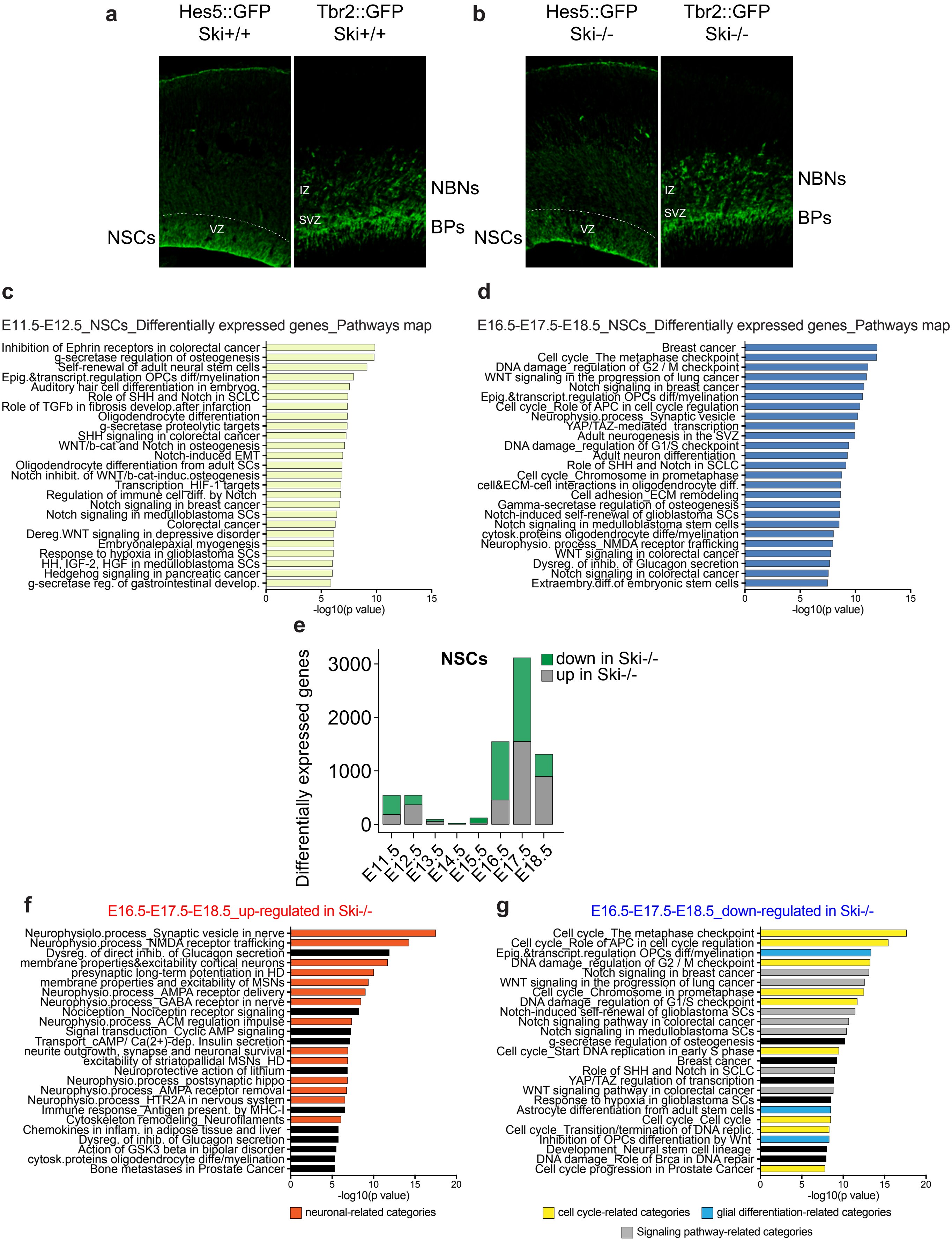

### ExData_Fig_2

#### Extended Data_Figure2

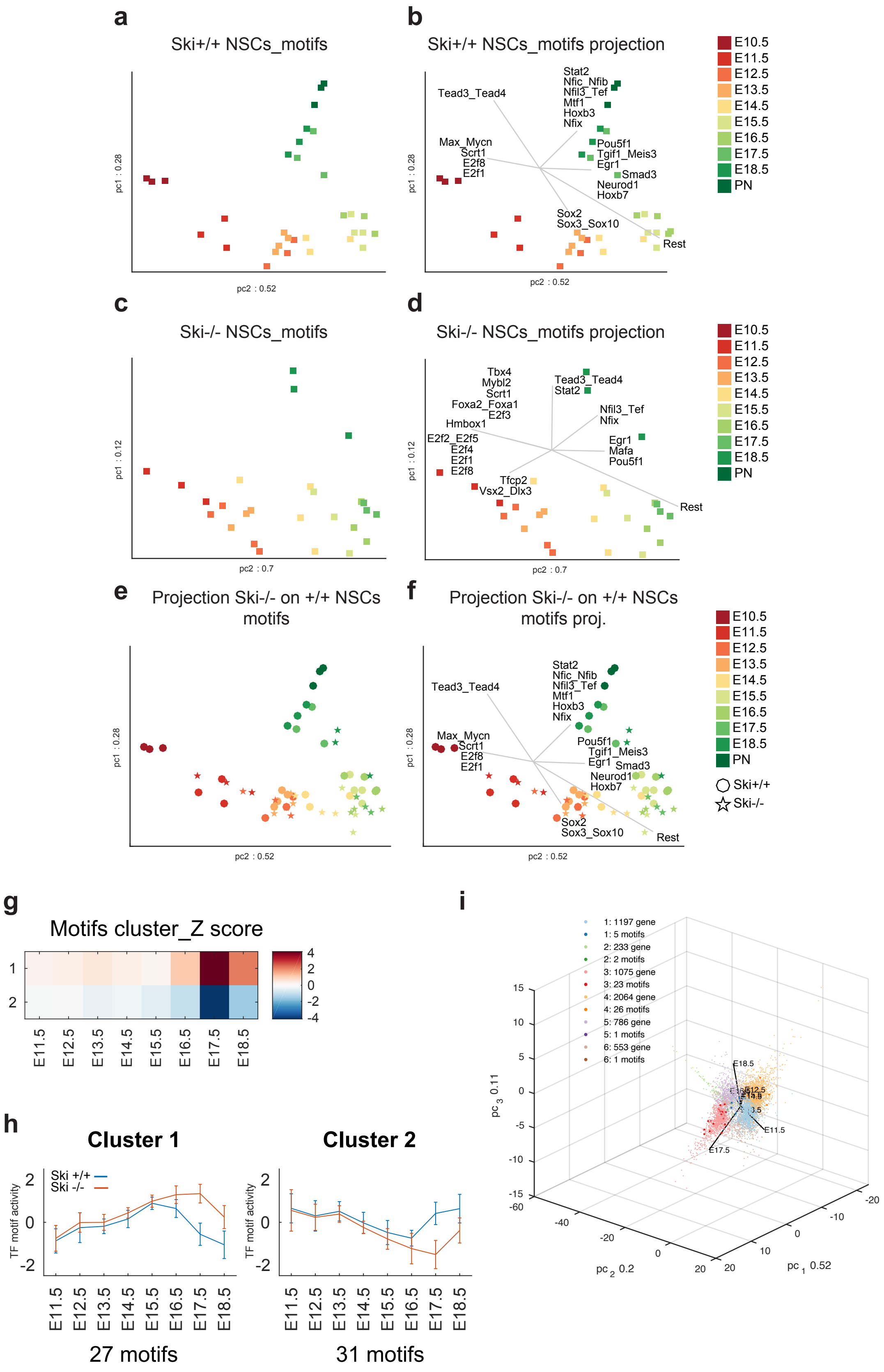

### ExData_Fig_3

Extended Data_Figure3

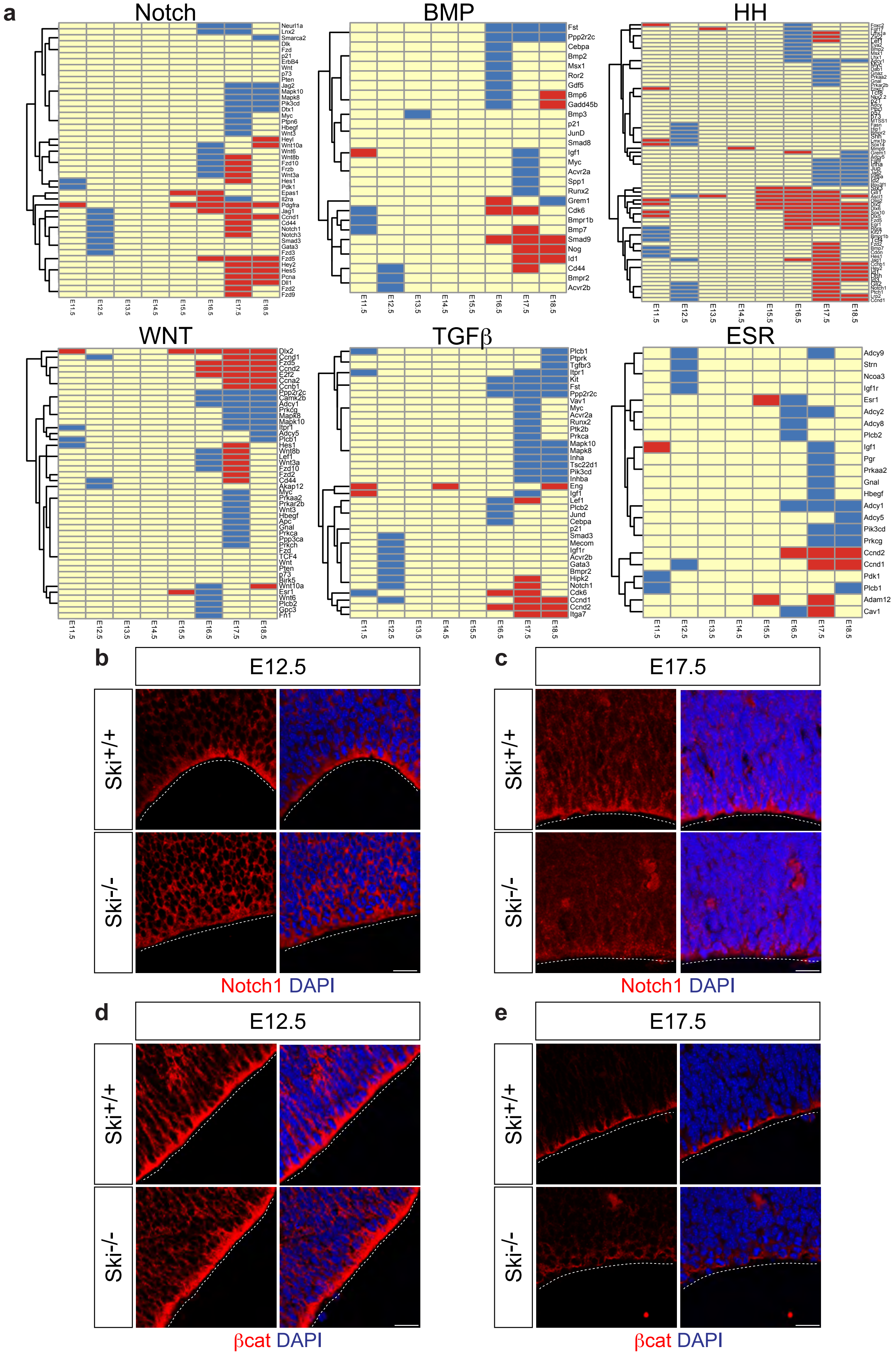

### ExData_Fig_4

#### Extended Data_Figure4

**a**

##### Neuronal enriched genes (Ye et al., 2014)

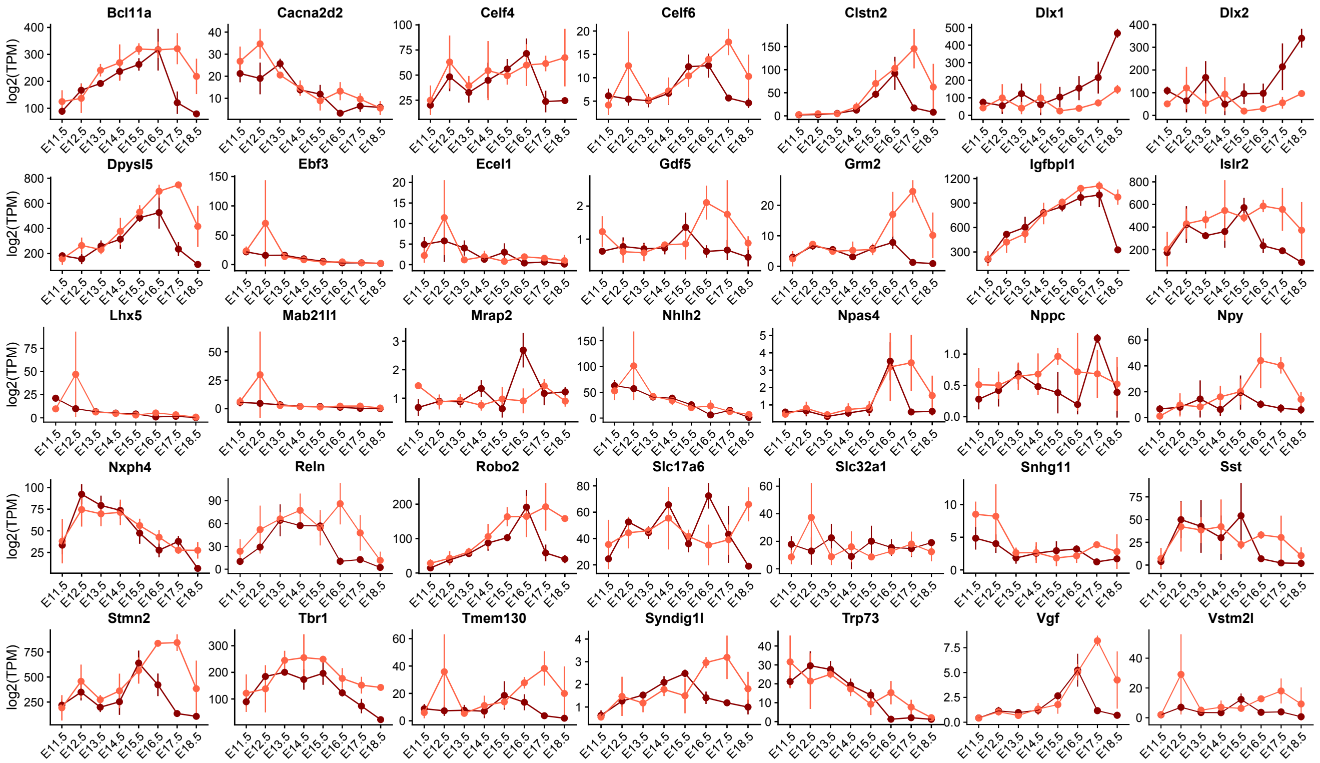


**b**

##### Glial enriched genes (Ye et al., 2014)

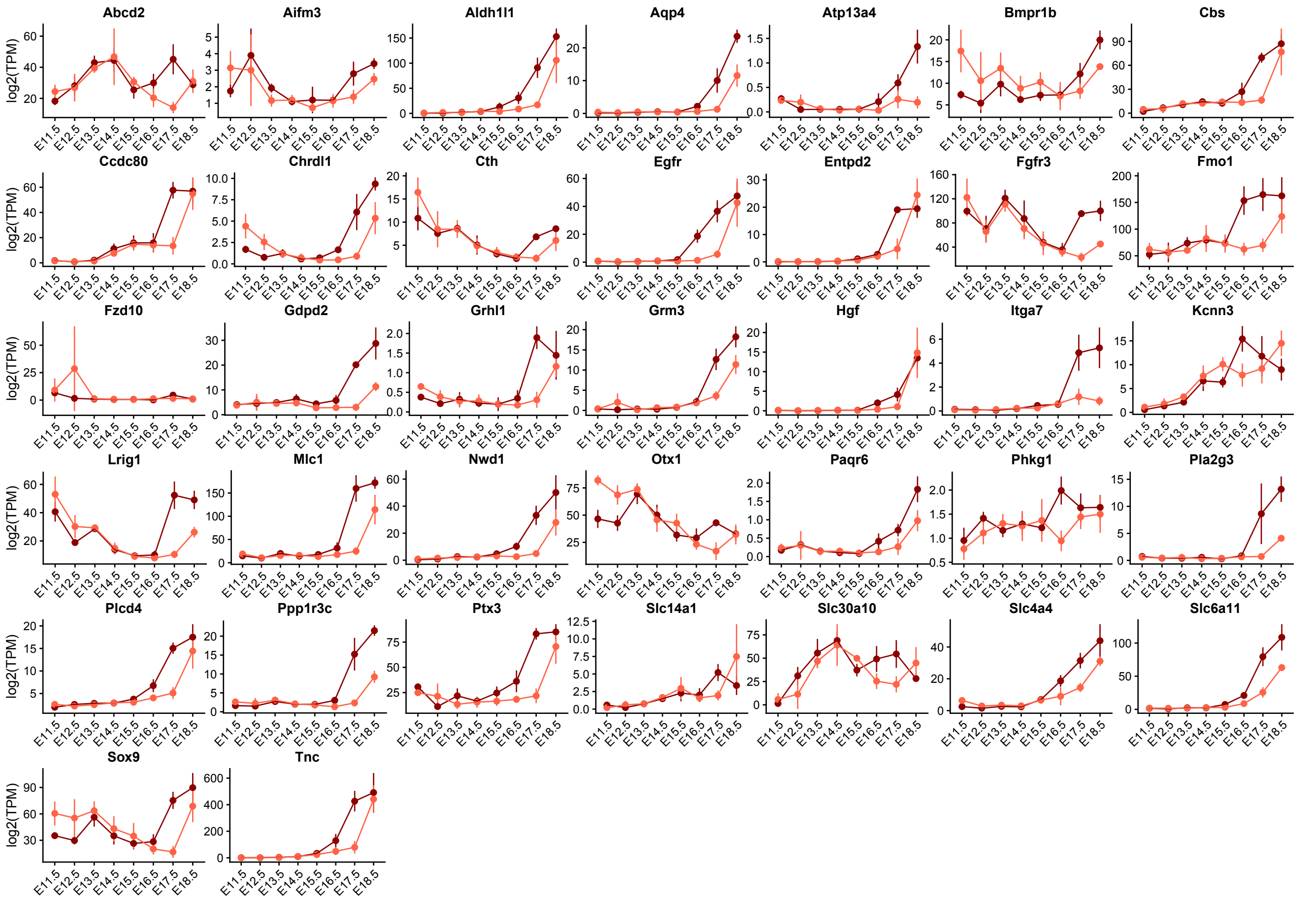

### ExData_Fig_5

Extended Data_Figure5

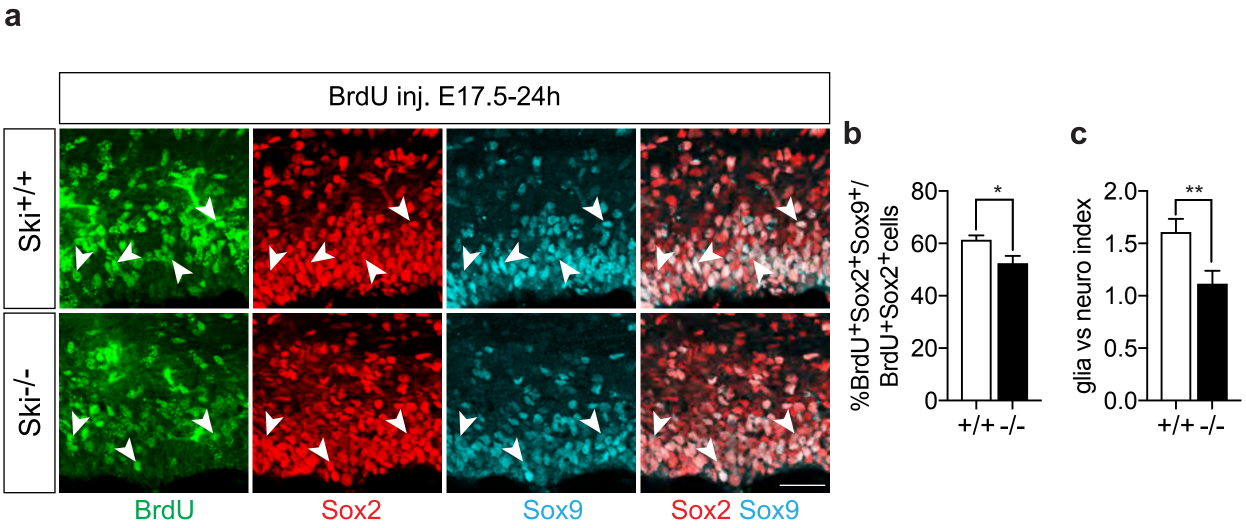

### ExData_Fig_6

Extended Data_Figure6

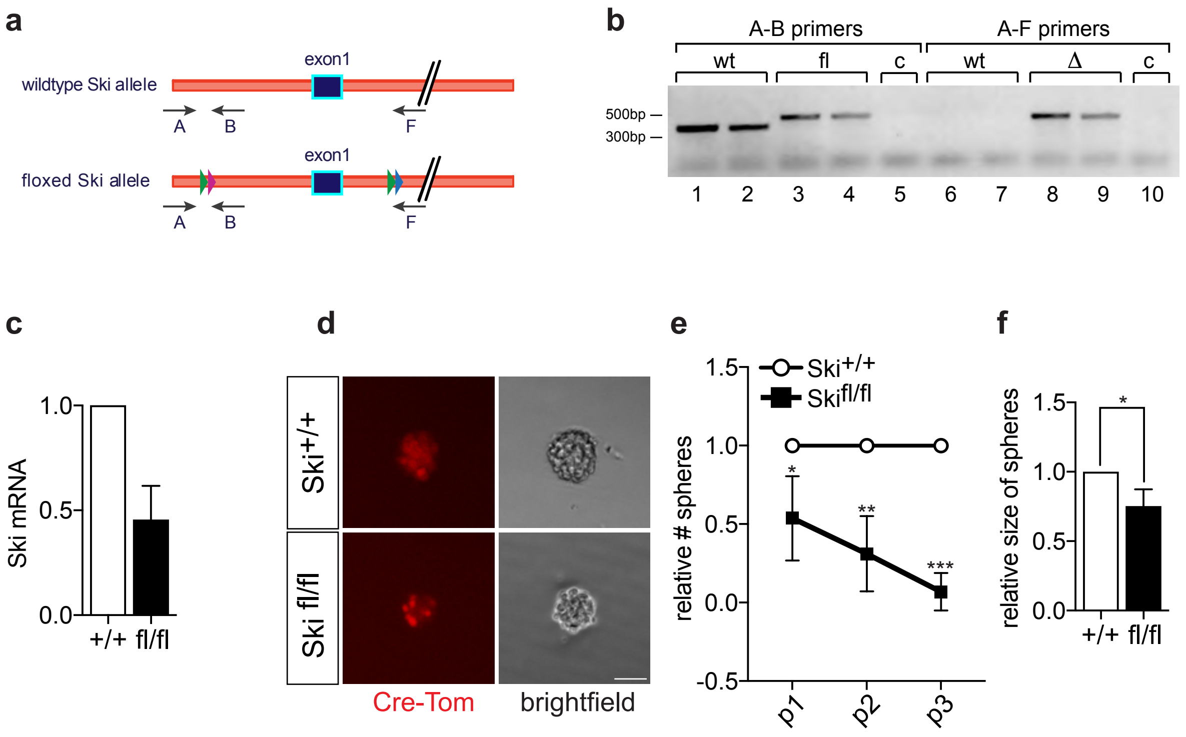
